# Multi-timescale reinforcement learning in the brain

**DOI:** 10.1101/2023.11.12.566754

**Authors:** Paul Masset, Pablo Tano, HyungGoo R. Kim, Athar N. Malik, Alexandre Pouget, Naoshige Uchida

## Abstract

To thrive in complex environments, animals and artificial agents must learn to act adaptively to maximize fitness and rewards. Such adaptive behavior can be learned through reinforcement learning^1^, a class of algorithms that has been successful at training artificial agents^2–6^ and at characterizing the firing of dopamine neurons in the midbrain^7–9^. In classical reinforcement learning, agents discount future rewards exponentially according to a single time scale, controlled by the discount factor. Here, we explore the presence of multiple timescales in biological reinforcement learning. We first show that reinforcement agents learning at a multitude of timescales possess distinct computational benefits. Next, we report that dopamine neurons in mice performing two behavioral tasks encode reward prediction error with a diversity of discount time constants. Our model explains the heterogeneity of temporal discounting in both cue-evoked transient responses and slower timescale fluctuations known as dopamine ramps. Crucially, the measured discount factor of individual neurons is correlated across the two tasks suggesting that it is a cell-specific property. Together, our results provide a new paradigm to understand functional heterogeneity in dopamine neurons, a mechanistic basis for the empirical observation that humans and animals use non-exponential discounts in many situations ^10–14^, and open new avenues for the design of more efficient reinforcement learning algorithms.

## Main

The ability to anticipate forthcoming events is crucial in choosing the right course of action. Predictive models have been a primary contender for the function of the cortex ^15,16^ and are at the core of recent proposals to design intelligent artificial systems ^17,18^. Many of these proposals rely on temporal difference (TD) reinforcement learning (RL) in which the TD learning rule is used to learn predictive information ^1,19^. By updating current estimates based on future expected estimates – TD methods have been remarkably successful in solving tasks that require predicting future rewards and planning actions to obtain them ^2,20–23^. In parallel, the TD learning rule has been used to explain the activity patterns of dopamine neurons in the midbrain, one of the classic examples where a normative computation has been successfully assigned to a genetically defined neuron type ^7–9^. However, there is mounting evidence suggesting that the representations encoded in dopamine neurons are far richer and more complex than a simple scalar reward prediction error ^24–32^, prompting reconsideration of the computational framework.

The standard formulation of TD learning assumes a fixed discount factor (that is, a single learning timescale) which, after convergence, results in exponential discounting: the value of a future reward is reduced by a fixed fraction per unit time (or time step). Although this formulation is important for simplicity and self-consistency of the learning rule, it is well known that humans and other animals do not exhibit exponential discounting when faced with inter-temporal choices. Instead, they tend to show hyperbolic discounting: there is a fast drop in value followed by a slower rate for further delays^10,12,33^. Far from being irrational, non-exponential discounting can be optimal depending on the uncertainty in the environment as has been documented in the behavioral economics and foraging literature ^13,14,34,35^. Humans and animals can modulate their discounting function to adapt to the temporal statistics of the environment and maladaptive behavior can be a signature of mental state or disease ^36–39^.

The TD rule can potentially be extended to learn more complex predictive representations than the mean discounted future reward of the traditional value function, both in artificial ^40–44^ and biological neural systems ^25,45,46^. A growing body of evidence points to the rich nature of temporal representations in biological systems ^47–49^ and particularly in the basal ganglia ^50–53^. Understanding how these rich temporal representations are learned remains a key question in neuroscience and psychology. An important component across most temporal-learning proposals is the presence of multiple timescales ^46,54–59^ which enables capturing temporal dependencies across a diverse range of durations: shorter timescales typically handle rapid changes and immediate dependencies, while longer timescales capture slow-changing features or long-term dependencies ^57^. Furthermore, work in AI suggests that the performance of deep RL algorithms can be improved by incorporating learning at multiple timescales ^60,61^. We therefore ask whether reinforcement learning in the brain exhibits such multi-timescale properties.

We first investigate the computational implications of multi-timescale RL. We then show that dopamine neurons encode predictions at diverse timescales, providing a potential neural substrate for multi-timescale reinforcement learning in the brain.

### Computational advantages of multi-timescale learning

We first examine the computational advantages of RL agents employing multiple timescales over those utilizing a single timescale. We start with a simple example environment where a cue predicts a future reward at a specific time (Fig. 1, see Methods). In standard RL algorithms, the agent learns to predict future rewards, compressed into a single scalar value, i.e. the sum of discounted future rewards expected from the current state ^1,19^: 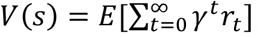, where *V*(*s*) is the value of the state *s, r*_*t*_ is reward at time *t*, and *γ* is the discount factor (0 < *γ* < 1, see Methods). *E* denotes the expectation over stochasticity in the environment and actions. Let *V*_*i*_ be the value learned using a discount *γ*_*i*_. Moving the discount factor *γ* out of the expectation, this equation can be rewritten (truncating at *t* = *T*) as

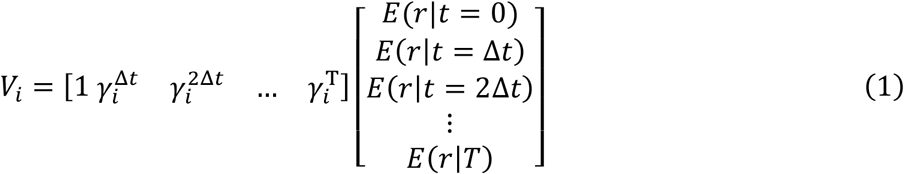

Where we assume that timesteps transitions are discrete and of size Δ*t* (see Methods). Thus, single-timescale learning projects all the timestep-specific expected rewards (*E*(*r*|*t*)) onto a single scalar (*V*_*i*_) through exponential discounting (Fig. 1a) and therefore entangles reward timing and reward size. When learning with multiple timescales, instead of collapsing all future rewards onto a single scalar, there is vector of value predictions, each computing value with its own discount factor *γ*_*i*_^45^:

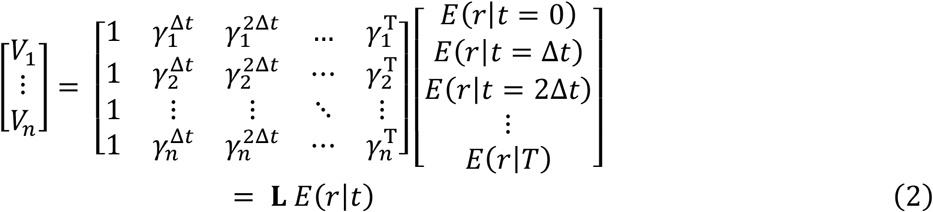

The last equality shows that the array of values learned with multiple discounts (Value space in Fig. 1b) corresponds to the Z-transform (i.e., the discrete Laplace transform) of the array that indicates the expected reward at all future timesteps (Temporal space in Fig. 1b). Since the Z-transform is invertible, the agent employing TD learning with multiple timescales can decode the expected temporal evolution of rewards from the representation of values that it learned, by applying a fixed, regularized decoder *L*^−1^ to the learned values ^45,62^ (Fig. 1b, fourth panel illustrates a situation with one reward per trajectory but this approach also work for multiple reward, see Methods and ref [^45^]). Intuitively, when learning with multiple timescales, the relative amplitude of the learned cue values as a function of discount factor (Value space in Fig. 1b) depends only on reward timing, and thus the agent can decode reward timing independently of reward magnitude.

**Figure 1.**
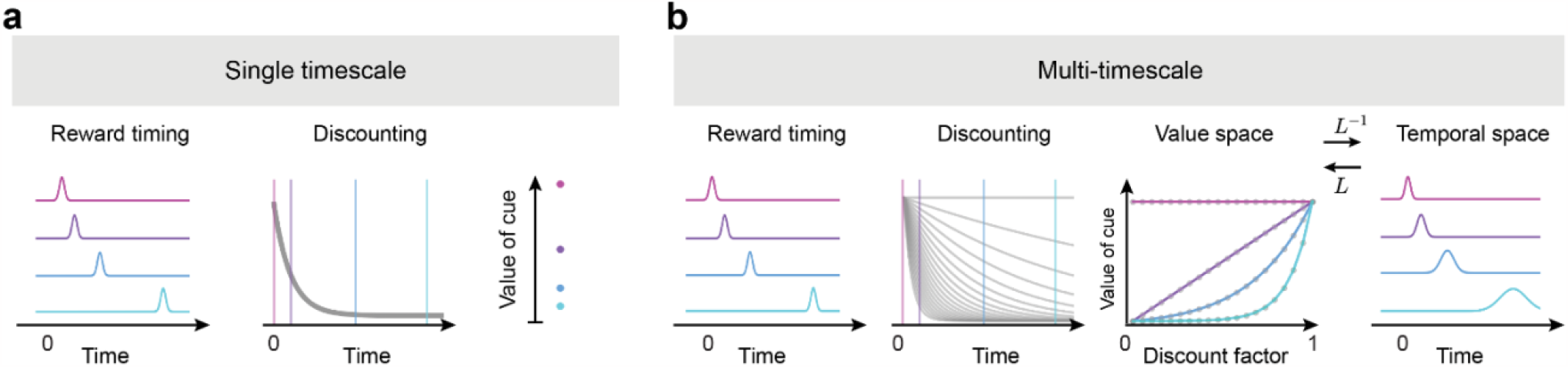
Single timescale and multi-timescale reinforcement learning. **a**, In single-timescale value learning, the value of a cue (at t = 0) predicting future rewards (first panel) is evaluated by discounting these rewards with a single exponential discounting function (second panel). The expected reward size and timing are encoded, but confounded, in the value of the cue (third panel). **b**, In multi-timescale value learning, the same reward delays are evaluated with multiple discounting functions (second panel). The relative value of a cue as a function of the discount depends on the reward delay (third panel). A simple linear decoder based on the Laplace transform can thus reconstruct both the expected timing and magnitude of rewards (fourth panel).

To illustrate the computational advantages of Laplace-transform multi-timescale agents, we consider several simple example tasks. The agent navigates through a linear track (a sequence of 15 states), where it encounters a reward of a certain magnitude (*R*) at a specific time point (*t*_*R*_, see Fig. 2a). The value of *R* and *t*_*R*_ changes across episodes and remains constant within episodes. Each episode is initiated by a cue presented at the initial state (*s*). Within each episode, the agent first learns the expected future rewards (i.e. the value, *V*_*γ*_(*s*)) predicted by the cue using a simple RL algorithm (*N* backups of tabular TD learning) employing one or multiple discount factors. Using the learned values associated with the cue, the agent then performs various tasks, using a deep neural network (DNN) trained across episodes with a policy gradient [PG] method; Fig. 2b and see Methods for details). Therefore, in our model, multi-timescale values are not used directly to produce behavior. Instead, they act as an enriched state representation from which task-specific behavior can be subsequently decoded (similarly to actor-critic and representation learning architectures like distributional RL ^41^). Our goal is to evaluate the advantages of the multi-timescale value representation over the single-timescale one.

**Fig 2.**
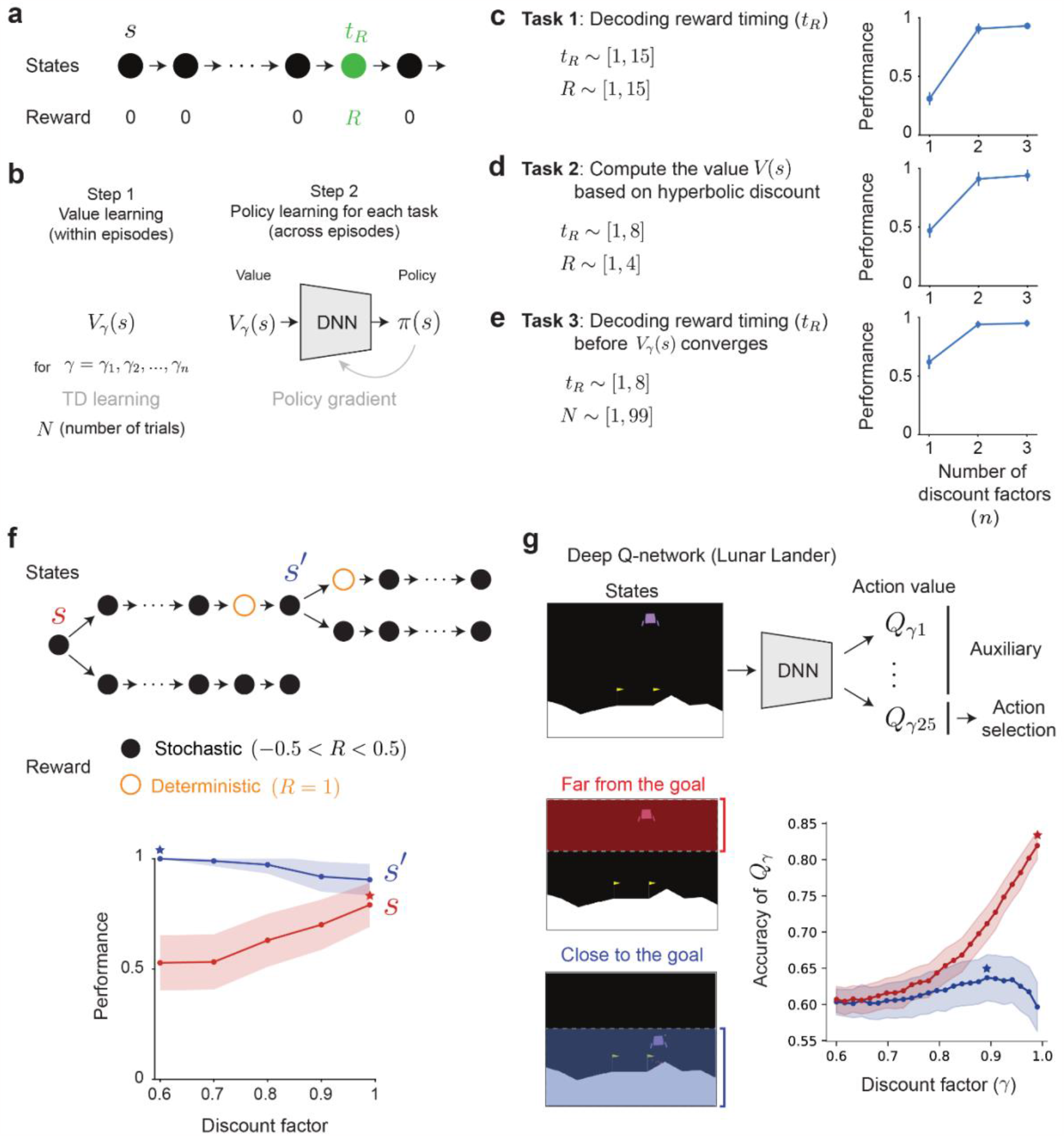
Computational advantages of multi-timescale reinforcement learning. **a**, Experiment to compare single-vs. multi-timescale learning. **b**, Architecture to evaluate multi-timescale advantages. In each episode (defined by a specific R, t_R_ and N) the value function is learned via tabular updates. The policy gradient network is trained across episodes to maximize the accuracy of the report. **c**, The timing t_R_ and reward size R is varied across episodes, the task of the policy gradient (PG) network is to report t_R_ . **d**, The timing t_R_ and reward size R is varied across episodes, the task is to report the inferred value of s using a hyperbolic discount. **e**, The timing t_R_ and number of sampled trajectories N is varied across episodes, the task of the policy gradient (PG) network is to report t_R_ . In **c-e**, Performance is reported after 1,000 training episodes. Error bars are the standard deviations (s.d.) across 100 test episodes and 3 trained policy gradient (PG) networks. **f**, Myopic learning bias. Top: Task structure to evaluate the learning bias induced by the discount factor, the three dots collapse 5 transitions between black states. Bottom: Performance at selecting the branch with the large deterministic reward under incomplete learning conditions. At state s (orange), agents with larger discount factors (far-sighted) are more accurate. At state s’ (blue), agents with a small discount factor (myopic) are more accurate. Error bars are half s.d. across 10,000 episodes, maximums are highlighted with stars. **g**, Top: Architecture that learns about multiple timescales as auxiliary tasks. Bottom: Accuracy of the Q-values in the Lunar Lander environment as a function of their discount factor, estimated as the fraction of concordant state pairs between the empirical value function and the discount specific Q-value estimated by the network, when the agent is close to the goal (blue) or far from the goal (orange), see Methods for details. Error bars are s.e.m across 10 trained networks, maximums are highlighted with stars.

#### Task 1: disentangling reward timing and reward magnitude

We first asked whether an agent can correctly discern the magnitude (*R*) and the timing (*t*_*R*_) of reward separately (Fig. 2c). We vary *R* and *t*_*R*_ across episodes. In each episode, the agent learns the values of states using 1, 2 or 3 discount factors. We then train the DNN across episodes to decode the timing of the reward (*t*_*R*_) with the vector of values associated with the cue {*V*_*γ*_(*s*)} as its input. With a single timescale, perfect performance is unattainable: a high value at the cue could signify a small reward in the near future or a large reward in the distant future. In contrast, the pattern of values across discount factors (third panel in Fig. 1b) is invariant to reward magnitude. As a result, multi-timescale agents can disentangle the timing (*t*_*R*_) and the magnitude (*R*) of reward (Fig. 2c, right, Extended Data Fig. 1a-c). Generally, the precision at which the timing and magnitude can be recovered depends on the number of discount factors being used (Extended Data Fig. 1a-c,j-l).

#### Task 2: learning values with non-exponential temporal discounts

While several tasks can be optimally solved by knowing the exponentially discounted state-values (i.e., where the value of a reward at time *t* decreases as *γ*^*t*^), the optimal temporal discount in a specific task depends on its temporal contingencies like its hazard rate, the cost of time and the uncertainty over time ^14,60^. Indeed, human and animal judgements are generally more consistent with a hyperbolic discount (i.e., decreasing as 1/(1+*γt*)) than an exponential one ^10,12,33^. However, the bootstrapping process of traditional TD value learning naturally converges to exponentially discounted values, so to perform optimally across tasks with arbitrary temporal contingencies, TD-learning agents need to adapt their exponentially discounted values to arbitrary, possibly non-exponential discounts. Crucially, multi-timescale systems encode the expected reward magnitudes at all future times (*E*[*r*|*t* = 0], *E*[*r*|*t* = Δ*t*], *E*[*r*|*t* = 2Δ*t*], …) in the inverse temporal Laplace space (i.e., after transforming the multi-timescale value estimates with *L*^−1^, see Fig. 1b). Consequently, they could weight the time-specific expected rewards with any chosen discount weights (e.g.*w*_0_*E*[*r*|*t* = 0] + *w*_1_*E*[*r*|*t* = Δ*t*] + ⋯) to retrieve the specific discount necessitated by the task. We demonstrate this in a task where the agent goal is to report the value of the initial state (*s*) using a hyperbolic discount (i.e. the value of a reward *R* at time *t*_*R*_ is *R* / (1+0.9*t*_*R*_)). With a single timescale, the learned exponentially discounted value cannot be accurately adapted into a hyperbolic one, but multi-timescale systems can reliably report the hyperbolic value of the cue given a diversity of exponential ones (Fig. 2d, Extended Data Fig. 1d-f, see Methods).

#### Task 3: inferring temporal information before convergence

In the above example (Fig. 2c), we showed that multi-timescale agents can disentangle the timing and the magnitude of rewards, which are typically intertwined in agents that rely on a single discount factor. This occurs because the shape of value function across discount factors encodes the proximity to rewards (Fig. 1b, third panel). We further hypothesized that, multi-timescale agents can leverage this advantage of extracting timing information even before value learning has fully converged. Consider an agent that has encountered a reward only a limited number of times (*N*). For single-timescale systems, a high value of the cue could be due to a short delay (*t*_*R*_) or simply because the value estimate has undergone more positive updates from an initial value of 0. In contrast, the shape of values encoded *across* discount factors is invariant to the number of reward encounters (*N*), to the extent that all value estimates depart from similar baselines and share similar learning parameters. As a result, multi-timescale agents can decode the time of reward (*t*_*R*_) even in situations where learning is incomplete (Fig. 2e, Extended Data Figs. 1g-i and 2, see Methods).

#### Task 4: state-dependent discount factor

Moreover, multi-timescale systems can preferentially adjust between myopic and farsighted perspectives based on the present circumstances. Consider a slightly more intricate maze with two branching points (Fig. 2f). In this maze, each state is associated with a random reward drawn uniformly between –0.5 and 0.5, except for two states (*s* and *s’*, orange circles) which result in a deterministic reward of 1. The optimal strategy in this scenario is to move upwards at both states *s* and s’, we define performance as the fraction of optimal choices across episodes. When learning from a limited number of experiences, the smaller stochastic rewards can overpower the larger deterministic rewards, making it challenging to achieve optimal performance. At state *s*, only far-sighted agents can discern the significance of the large deterministic rewards, thereby causing myopic agents to perform near chance at *s*. At state *s*’, the situation is reversed. Far-sighted agents not only integrate the close-by large reward but also all the stochastic rewards farther in the future. Myopic agents, in contrast, assign greater weight to the reward of 1 compared to the future stochastic rewards, thus enabling optimal performance at *s’*. Therefore, only agents that could dynamically adapt between being far-sighted at *s* and myopic at *s’* can attain optimal performance when learning from limited experiences. Indeed, the multi-timescale of Fig. 2b achieves in this task a maximum performance of 83±1% with a single discount and a performance of 94±1% with two discounts. The superior performance is due to its demonstrated ability to discern the temporal distance to the relevant events in the environment (here, the large deterministic rewards), and subsequently focus on the myopic or far-sighted values depending on the estimated distance. We also observe the benefits of the myopic learning bias in more realistic navigation scenarios (Extended Data Figs. 1m-o and 3) as well as in more complex Deep RL settings where additional timescales act as auxiliary tasks (Fig. 2g, see Methods).

To summarize, in multi-timescale value systems the vectorized learning signal robustly contains temporal information independently of the information about reward magnitude. This property empowers agents to selectively focus on either myopic or far-sighted estimates depending on the current situation.

### The diversity of discount factors across dopamine neurons conveys distributional information about the timing of future rewards

In the previous section, we demonstrated the computational advantages of learning with multiple discount factors for an RL agent. Building upon these findings, we next investigated whether the brain employs such multi-timescale RL. Toward this goal, we examined the activity of dopamine neurons, which are believed to encode the TD error term in RL algorithms.

To characterize the discounting properties of individual dopaminergic neurons, mice were trained in a cued delay task ^50,63^ in which on a given trial, one out of four distinct odor cues indicated its associated timing of a water reward (Fig. 3a). These odor cues were preceded by a trial start cue (green computer screen) by 1.25s. The trial start cue reduced the timing uncertainty of the odor cue and ensured that the responses of dopaminergic neurons to the odor cues were mostly driven by a valuation signal rather than a saliency signal ^64,65^. Mice showed anticipatory licking prior to reward delivery. Importantly, the onset of the anticipatory licking was delayed for trials with cues predicting longer reward delays, indicating that the mice learned the delay contingencies (Fig. 3b). We recorded optogenetically identified single dopamine neurons in the ventral tegmental area (VTA) (*n* = 78, see Methods). We focused our analysis on neurons (*n* = 50) who passed the selection criteria (including mean cue response firing rate above 2 spikes/s, positive goodness of fit on test data, see Methods). As expected from RL theory and the prediction error framework, the average responses to the reward cue decreased as the predicted reward timing increased ^50,63^(Fig. 3c, Extended Data Fig. 4a-b). However, cue responses of individual neurons showed a great diversity of discounting across the reward delays ranging from neurons responding strongly only to the cue indicating the shortest delay to neurons with a gradual decay of their response with cued reward delay (Fig. 3d-e).

**Figure 3.**
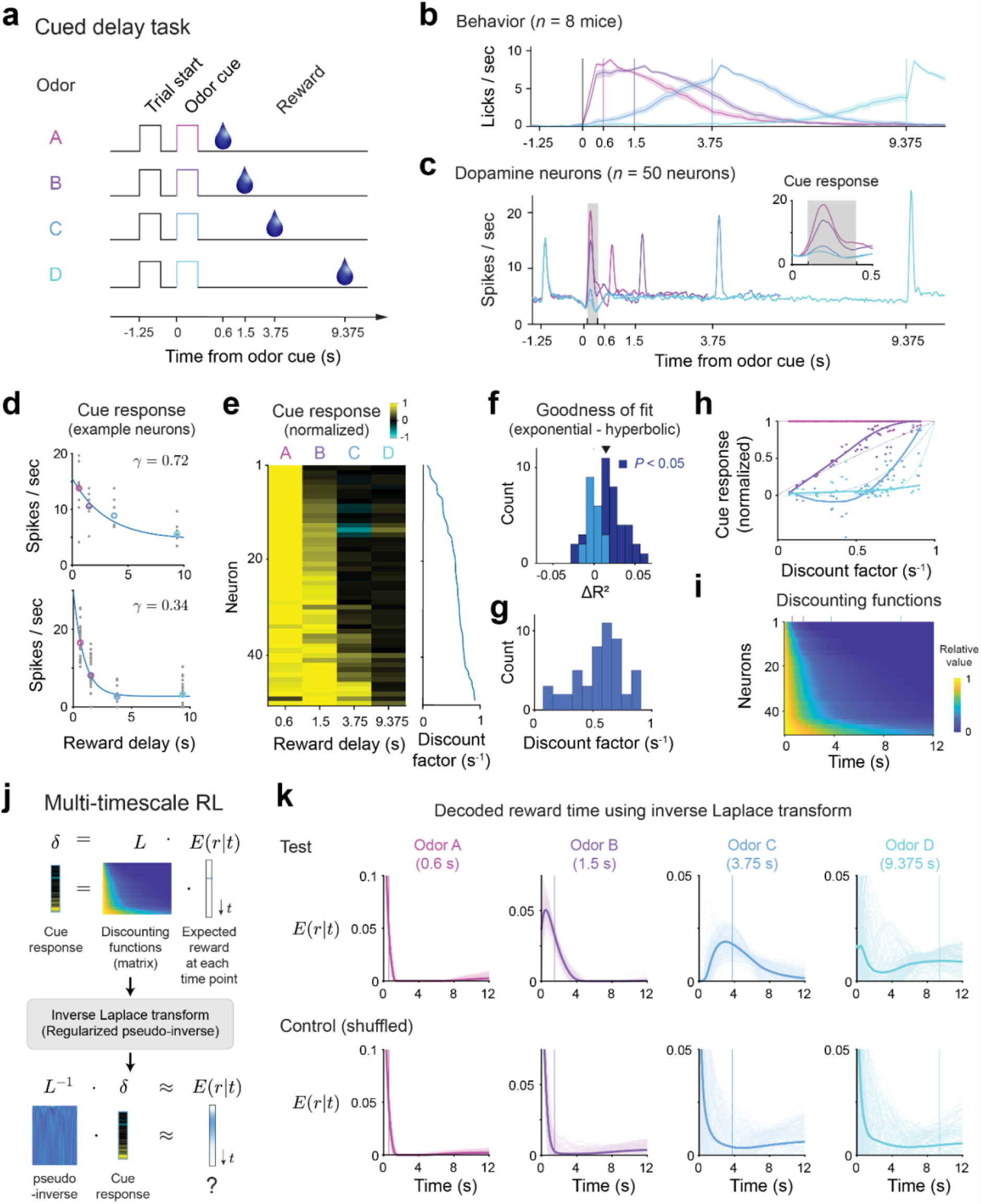
Dopamine neurons exhibit a diversity of discount factors that enables decoding of reward delays. **a**, Outline of the task structure. **b**, The mice exhibit anticipatory licking prior to reward delivery for all 4 reward delays indicating that they have learned task contingencies (mean across behavior for all recorded neurons, shaded error bar indicates 95% confidence interval using bootstrap). **c**, Average PSTH across the task for the 4 trial types. Inset shows the firing rate in the 0.5s following the cue predicting reward delay. The firing rate in the shaded grey box (0.1s < t < 0.4s) was used as the cue response in subsequent analysis. **d**, Example of fits of the responses to the cue predicting reward delay of two single neurons with high (top panel) and low (bottom panel) discount factors. **e**, Normalized response to the cues predicting reward delays across the population. For each neuron, the response was normalized to the highest response across the 4 possible delays. Inset on right, corresponding inferred discount factor for each neuron. **f**. The exponential model is a better fit to the data than the hyperbolic one as quantified by distance of mean R^2^ to the unit line. Mean = 0.0147, P = 2.2 x 10^-5^, two-tailed t-test. Shading indicated significance for single neurons across bootstraps (dark blue: P < 0.05). **g**, Distribution of inferred discount factors across neurons. For each neuron, the discount factor was taken as the mean discount factor across bootstraps. **h**. Shape of the relative population response as a function of reward delay. Normalized to the strongest cue response for each neuron. Thick lines, smoothed fit, dotted lines, theory, dots, responses of individual neurons. **i**, Discount matrix. For each neuron we plot the relative value of future events given its inferred discount factor. Neurons are sorted as in panel **d** by increasing inferred value of the discount factor. Vertical bars on top of panel are color coded to indicate timing of the rewards in the task. **j**, Outline of the decoding procedure. We compute the singular value decomposition (SVD) of the discount matrix **L**. Then, we use the SVD to compute a regularized pseudo-inverse **L**^-1^. Finally, we normalize the resulting prediction into a probability distribution. **k**, The subjective expected timing of future reward E(r|t) can be decoded from the population responses to the cue predicting reward delay. Decoding based on mean cue responses for test data (top row, see Methods). The ability to decode the timing of expected future reward is not due to a general property of the discounting matrix and collapses if we randomize the identity of the cue responses (bottom row, see Extended Data Fig. 5e and Methods).

To characterize the discount properties of individual neurons, we fit them individually using both an exponential discount model and a hyperbolic discount model. The exponential model provided a better fit to the neurons’ responses than the hyperbolic model (*P* = 2.2 x 10^-5^, two-tailed *t*-test; Fig. 3f and Extended Data Fig. 4c-e, see Methods) contrary to a previous observation in non-human primates ^63^. Organism level hyperbolic-like discounting can, therefore, arise from the diversity of exponential discounting in single neurons, as discussed above with artificial agents (Fig. 2d, see also refs [^14,55,60^]). This view is consistent with the wide distribution of inferred discount factors obtained across the population (0.56 ± 0.21 s^-1^, mean ± s.d., Fig. 3g). Fits to simulated data suggest that our estimate of inferred parameters is robust and primarily constrained by the number of trials (Extended Data Fig. 4f-h, see Methods).

As we have shown above, artificial agents equipped with diverse discount factors exhibit various advantages. One key aspect contributing to these advantages is their unique ability to independently extract reward timing information, which is lacking in single timescale agents. We next asked whether dopamine neurons provide a population code in which the structured heterogeneity across the population enables decoding of reward timing or the expected reward across time, *E*(*r*|*t*). Mathematically, this transformation can be achieved by the inverse Laplace transform (or its discrete equivalent the Z-transform, Fig. 3j) ^45,57,62^. In our data set, the dopaminergic cue responses for each reward delay exhibited unique shapes as a function of discount factors, suggesting that reward timing information is embedded in the dopaminergic population responses (Fig. 3h, compare with Fig. 1b, third panel). The temporal horizon across the population, which underlies these cue responses, can be visualized through the discount matrix which indicates for each neuron the relative value of a future reward depending on the inferred discount factor (Fig. 3i).

If the dopaminergic population code is consistent with the Laplace code explored above (Fig. 1-2), reward timing should be recoverable from the dopamine neurons’ cue responses with a regularized discrete inverse Laplace transform of the neural activity (which does not require training a decoder). In our task, we can use the TD-error driven cue responses (instead of the value in equation 2) as they are driven by the discounted future value (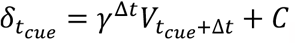, see Methods). This implies that the right-hand side of equation 2 can be approximated by the population dopamine responses. We used a pseudo-inverse of the discount matrix (computed using half of all trials) based on regularized singular value decomposition to approximate the inverse Laplace transform (Fig. 3j, Extended Data Fig. 5a-d, see Methods and ref[^45^]) and applied it to dopamine neuron cue responses (computed on the held out half of the trials). Remarkably, the decoder was able to predict reward timing, closely matching the true reward delay (Fig. 3k, top row). This prediction was lost if we shuffled the neuron identities indicating that it is not a generic property of the discount matrix (Fig. 3k, bottom row). We quantified this decoding by computing a distance metric (using 1-Wasserstein distance) between the true and predicted reward delay across conditions (*P* = 1.2 x 10^-4^ for 0.6 s reward delay, *P* < 1.0 x 10^-20^ for the other delays, one-tailed Wilcoxon signed rank test; Extended Data Fig. 5e, see Methods). Predictions from the model were more accurate than an alternative model with a single discount factor (*P*_*t* = 0.6s_ = 1, *P*_*t* = 1.5s_ < 1.0 x 10^-31^, *P*_*t* = 3.75s_ = 0.0135, *P*_*t* = 9.375s_ < 1.0 x 10^-14^, one-tailed Wilcoxon signed rank test; Extended Data Fig. 5f-g and see Methods). Consistent with the above observation that cue responses were fit better with exponential over hyperbolic discounting models, the accuracy of reward timing decoding was typically higher when using the discount matrix from the exponential model than the one from the hyperbolic model (*P*_*t* = 0.6s_ = 1, *P*_*t* = 1.5s_ < 1.0 x 10^-31^, *P*_*t* = 3.75s_ < 1.0 x 10^-33^, *P*_*t* = 9.375s_ < 1.0 x 10^-3^, one-tailed Wilcoxon signed rank test; Extended Data Fig. 6a-e). Furthermore, the decoding performance was comparable to simulated data with matched trial numbers, indicating that the remaining uncertainty in decoded reward timing is primarily driven by limited sample size in the data (e.g., the number of neurons and the number of trials per condition, Extended Data Fig. 6f-g and see Methods).

Together these results establish that dopamine neurons compute prediction errors with a heterogeneity of discount factors and show that the structure in this heterogeneity can be exploited by downstream circuits to decode reward timing.

### Heterogeneity of discount factors explains diverse ramping activity across dopamine neurons

In the task above (Fig. 3), prediction errors in dopamine neurons were measured through discrete transitions in the value functions at the time of cue. In more naturalistic environments, value might change more smoothly, for example when an animal approaches a goal ^66^. In these tasks, ramps in dopaminergic signaling have been initially interpreted as quantifying value functions ^32,66^ but were recently shown to conform to the predictions of the TD learning model. Specifically, these ramps can be understood as moment-by-moment changes in values or as TD error along an increasingly convex value function in which the derivative is also increasing ^67–69^. Here we show that some of this heterogeneity can be understood as evidence for multi-timescale RL across dopamine neurons.

We analyzed the activity of optogenetically identified dopamine neurons (*n* = 90, see Methods and ref [^68^]) while mice traversed along a linear track in virtual reality (VR). Although mice were free to locomote, their movements did not affect the dynamics of the scene (see Methods and ref [^68^] for details). At trial onset, a linear track appeared, the scene moved at continuous speed and reward was delivered around 7.35 seconds after motion onset (Fig. 4a). The slope of ramping across neurons was on average positive (Fig. 4b-c) but single neurons exhibited a diversity of ramping activity (Fig. 4c-e) ranging from monotonic upward and downward ramps to non-monotonic ramps.

**Figure 4.**
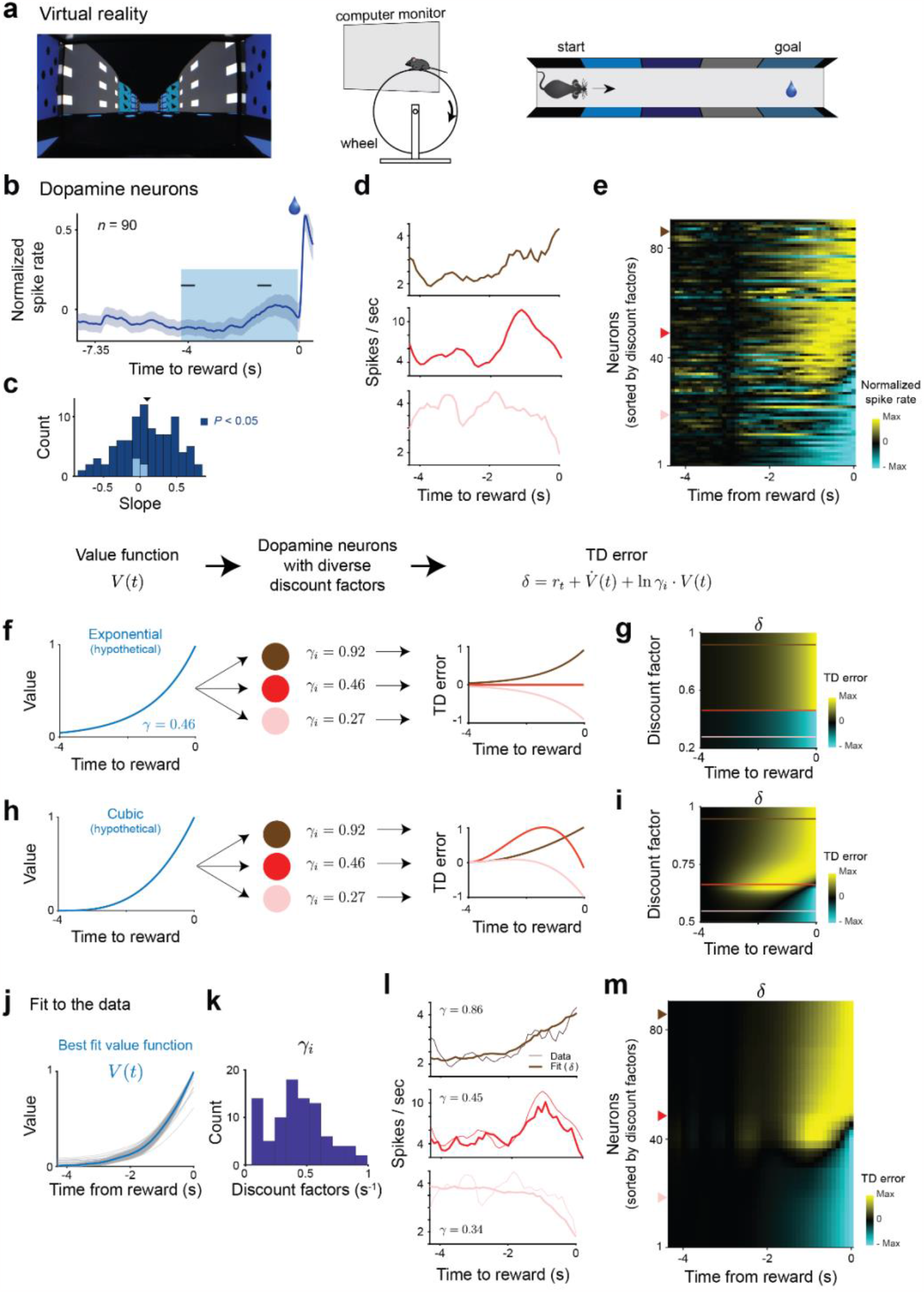
The diversity of discount factors across dopamine neurons explains qualitatively different ramping activity. **a**, Experimental setup. Left panel, View of the virtual reality corridor at movement initiation. Middle and right, Schematics of the experimental setup. **b**, Average activity of single dopaminergic neurons (n = 90) exhibit an upward ramp in the last few seconds of the track prior to reward delivery. **c**, The slope of the activity ramp (computed between the two black horizontal ticks in panel **b**) is positive on average but varies across neurons (population: mean slope = 0.097, P = 0.0175. Single neurons: positive and P < 0.05: n = 53; negative and P < 0.05: n = 32; P > 0.05: n = 5, two-tailed t-test). **d**, Example single neurons showing diverse ramping activity in the final approach to reward including, monotonic upwards (dark red), non-monotonic (red) and monotonic downwards (light red) ramps. **e**, Individual neurons across the population exhibit a spectrum of diversity in their ramping activity. Neurons are sorted according to inferred discount factor from the common value function model (panel **k**). **f**, Diversity of ramping with an exponential value function. There is no TD error for an agent with the same discount factor as the parameter of the value function (red line). The TD error ramps upwards (downwards) if the discount factor is larger (smaller), dark red and light red lines respectively. **g**, Diversity of ramping as a function of discount factor for an exponential value function. **h**, Diversity of ramping with cubic value function. Agents with large (small) discount factor experience a monotonic positive (negative) ramp in their TD error (dark red and light red lines respectively). Agents with intermediate discount factors experience non-monotonic ramps (red line). **i**, Diversity of ramping as a function of discount factor for an exponential value function. Unlike in the exponential value function case, no agent matches its discount to the value function at all the time steps. **j**, The inferred value function is convex. Thin grey lines represent the inferred value function for each bootstrap. Thick blue line represents mean over bootstraps. **k**, Histogram of inferred discount factors. 0.42 ± 0.23 (mean ± s.d.). **l**, Example model fits for the single neurons shown in panel **d. m**, The model captures the diversity of ramping activity across the population. Neurons are ordered by inferred discount factor as in panel **e**.

We hypothesized that this seemingly puzzling heterogeneity can be understood as a signature of multi timescale reinforcement learning. Considering that the value function is set by the limits on the precision of internal timing mechanisms and the reduction in uncertainty due to visual feedback ^69,70^, we first assume that heterogeneous dopamine neurons contribute to learning a common model of the environment and therefore share a common value function (see Methods). Depending on the shape of this value function, governed by the statistics of the environment being learned, the TD error from neurons with different discount factors will exhibit different type of activity ramps. At a given time, the sign of the TD error will depend on the relative scale of the upcoming increase in value and the reduction of this future value due to discounting. Given an increase in value *1/γ*_*o*_ (with *γ*_*o*_ < 1) a neuron with a discount factor smaller, equal or larger than *γ*_*o*_ *w*ill experience a negative, zero or positive TD error respectively (see Extended Data Fig. 7a and Methods). For an exponential value function (Fig. 4g, left panel), where the value increases by a fixed factor 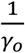 at every timestep, a neuron with discount factor *γ*_*o*_ will have no TD error during the entire visual scene (red line, Fig. 4f,g). A neuron with a higher (or lower) discount factor than *γ*_*o*_ will experience an upward (or downward) monotonic ramp in its activity (darker and lighter red line in Fig. 4f-g respectively). However, if the value function is non-exponential (for example cubic as a function of distance to reward, Fig. 4h, left panel), there will not be a neuron whose discount factor is able to match the increases in value function at all timesteps. Neurons with high or low discount factors will still ramp upwards or downwards (darker and lighter red line in Fig. 4h-i respectively), but neurons with intermediate discount factors will exhibit non-monotonic ramping (red line, Fig. 4h-i) as observed in the neural data.

To fit this model to the dopaminergic neurons, we used a bootstrapped constrained optimization procedure on a continuous formulation of the TD error ^69,71^ (see Methods) by fitting a non-parametric common value function and neuron-specific gains, baselines and discount factors. Although the gain and baseline activity scale the range of activity, only the interaction between the value function and the discount factor affects the shape of the TD error across time (see Methods). The heterogeneity of ramping activity across single neurons is explained (Fig. 4l-m) by a common convex value function (Fig. 4j) and a diversity of discount factors across single neurons (Fig. 4k). We did not observe a significant correlation between inferred parameters and the medio-lateral position of the implanted electrodes (Extended Data Fig. 7b-d). So far, we proposed a descriptive model with a common value function across neurons suggesting that single neurons predictions errors are pooled to create a single value function and world model. Recent models for distributed prediction errors across dopamine neurons have instead used parallel loops where individual neurons contribute to estimating sperate value functions ^25,45,72–75^. Instead of a common value function, the dopamine neurons can be part of independent loops and share a common expectation of reward timing. We obtained similar results in this common reward expectation model (see Methods and Extended Data Fig. 8).

Together these results show that diversity in slow changes in activity across single neuron (known as dopamine ramps) in environments with gradual changes in value can be explained by a diversity of discount factors and is a signature of multi-timescale reinforcement learning.

### Inferred discount factors for single neurons are correlated across the two behavioral tasks

Distributional RL and other distributed RL formulations provide agents with greater flexibility as they allow agents to adapt risk sensitivity and discounting to the statistics of the environment ^41,45,60,73^. However, they leave open the question of the biological implementation of this adaptivity. Specifically, the tuning of single dopamine neurons, controlled by the sensitivity to reward size or the discount factor, could be either a circuit property and therefore task and context specific or it could be a cell-specific property, with the contribution of different neurons recruited according to task demands. However, measurements of tuning diversity at the single neuron level are usually done in a single behavioral task ^25,28,76^, leaving open the question of this implementation across contexts.

Here, we characterized discount factors across two behavioral tasks and a subset (*n* = 43) of the single neurons analyzed above (Figures 3 and 4) were recorded on the same day in both behavioral tasks. Using this data set, we found that the discount factors inferred independently across the two behavioral tasks are correlated (Fig. 5a-b). Furthermore, in the cued delay task, we were able to decode subjective reward timing from population cue responses using the discount matrix built from the discount factors inferred in the virtual reality task (*P*_*t* = 0.6s_ = 1, *P*_*t* = 1.5s_ < 1.1 x 10^-20^, *P*_*t* = 3.75s_ < 3.8 x 10^-20^, *P*_*t* = 9.375s_ < 2.9 x 10^-5^, compared to shuffled data, Extended Data Fig. 9 and see Methods). These results suggest that the discount factor (or its ranking) is a cell-specific property and strongly constrains the biological implementation of multi-timescale reinforcement learning in the brain.

**Figure 5.**
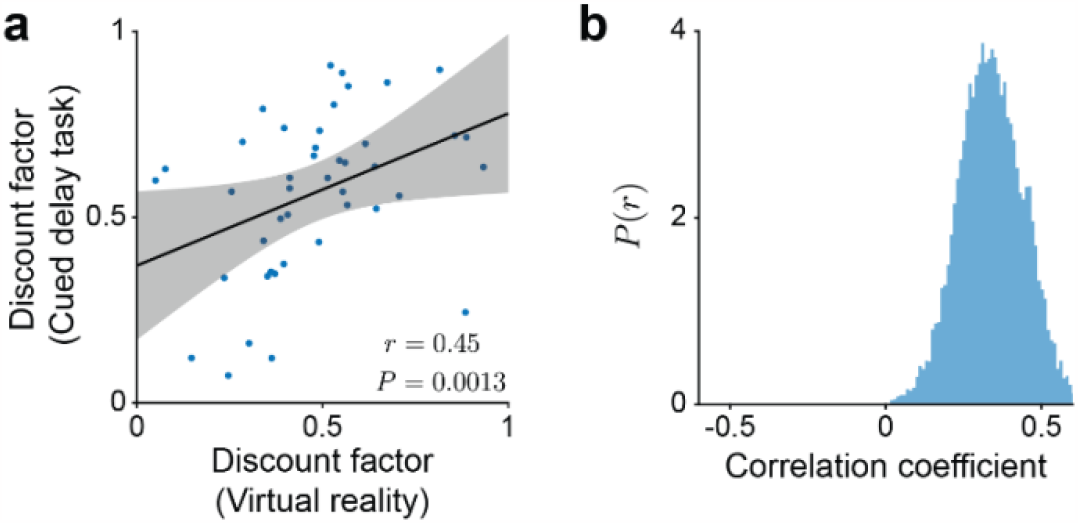
Discount factors of single dopaminergic neurons are correlated across behavioral contexts. **a**, Correlation between the discount factors inferred in the VR task and the discount factors inferred in the cued delay task (r = 0.45, P = 0.0013). **b**, Distribution of correlations between the discount factors across the two tasks for randomly sampled pairs of bootstrap estimates (0.34 ± 0.104, mean ± s.d., P < 1.0 x 10^-30^, two-tailed t-test).

## Discussion

In this work, we have analyzed the unique computational benefits of multi-timescale reinforcement learning agents and shown that we can explain multiple aspects of the activity of dopaminergic neurons through that lens.

The understanding of dopaminergic neurons as computing a reward prediction error from TD reinforcement learning algorithms has transformed our understanding of their function. However, recent experimental work expanding the anatomical locations of recordings and the task designs has shown heterogeneity in dopamine responses that is not readily explained within the canonical TD framework ^26,28,32,66,77,78^. However, a number of these seemingly anomalous findings can be reconciled and integrated within extensions of the RL framework, further reinforcing the power and versatility of the TD theory in capturing the intricacies of brain learning mechanisms ^24,25,29,45,69,72,74,75,79^. In this work, we reveal an additional source of dopaminergic heterogeneity: they encode prediction errors across multiple timescales. Together, these results indicate that at least some of the heterogeneity observed in dopamine responses reflects variations in key parameters within the RL framework. Thus, these results indicate that the dopamine system employs “parameterized vector prediction errors”, including a discrete Laplace transform of the future temporal evolution of the reward function, allowing for the learning and representation of richer information than what can be achieved with scalar prediction errors in the traditional RL framework.

The constraint on the anatomical implementation of multi-timescale RL suggested by the alignment of discount factors between the two tasks could also inform algorithm design. Adapting the discount factor has been used to improve performance in several algorithms, with proposed methods ranging from meta-learning an optimal discount factor ^80^, learning state dependent discount factors ^81,82^, or combining parallel exponentially discounting agents ^55,60,61^. Our results provide evidence supporting the third model but the recruitment mechanisms of the neurons to adapt the global discounting function with task or context and the link between anatomical location and discounting^53^ remain open questions. Similarly, the contribution of this vectorized error signal on the downstream temporal representations^49,51^ remains to be explored.

Understanding how this recruitment occurs will be a key step towards a mechanistic understanding of the contribution of this timescale diversity to calibration and miscalibration in intertemporal choices. There has been a conundrum that RL theories use exponential discounting while humans and animals often exhibit hyperbolic discounting. A previous study, that examined discounting in dopamine neurons, argued that single dopamine neurons exhibit hyperbolic discounting ^63^. However, they used uncued reward responses for zero reward delay, likely biasing the estimate toward hyperbolic (as responses to unpredicted rewards are typically large and potentially contaminated by salience signals). In contrast, our data are consistent with exponential discounting at the level of single neurons, suggesting that RL machinery defined by each dopamine neuron conforms to the rules of a simple RL algorithm. Hyperbolic-like discounting can occur when these diverse exponential discounting are combined at the organism level ^14,36,55^. More generally, the relative contribution of multiple timescales to the global computation governs the discount function at the organism level and should be calibrated to the uncertainty in the hazard rate of the environment ^14^.

Appropriately recruiting the heterogeneity of discount factors is therefore important to adapt to the temporal uncertainty of the environment. This view draws parallels with the distributional RL hypothesis that naturally fits with current work on anhedonia as a miscalibration of optimism and pessimism can lead to biases in the learned value ^25^. Miscalibration of the discounting spectrum can lead to excessive patience or impulsivity. A bias in this distribution due to genetical, developmental or transcriptional factors could bias the learning at the level of the organism towards short- or long-term goals. Behaviorally such bias would manifest itself as an apparent impulsivity or lack of motivation, leading to a potential mechanistic interpretation of these maladaptive behaviors. Similarly, this view could guide the design of algorithms that recruit and leverage these adaptive temporal predictions.

Our study establishes a new paradigm to understand the functional role of prediction error computation in dopaminergic neurons and opens new avenues to develop mechanistic explanations for deficits in intertemporal choice in disease and inspire the design of new algorithms.

## Methods

### Animal care and surgical procedures

The mouse behavioral and electrophysiological data presented here was collected as part of a previous study where all experimental procedures are described in details ^68^. As described in this study, all procedures were performed in accordance with the National Institutes of Health Guide for the Care and Use of Laboratory Animals and approved by the Harvard Animal Care and Use Committee.

We used a total of 13 adult C57/BL6J DAT-Cre male mice. Mice were backcrossed for over 5 generations with C57/BL6J mice, Animals were singly housed after surgery on a reverse 12 hr dark/12 hr light cycle (dark from 7am to 7pm). Single dopaminergic neurons were optogenetically identified using custom built micro drives with 8 tetrodes and an optical fiber as described in our previous study ^68^. Significance was assessed using the stimulus associated spike latency test (SALT) ^83^.

All mice (*n* = 13) were used in the virtual reality task and 8 of those were also used in the cued delay task. The targeted medio-lateral (ML) location varied from 320μm to 1048μm for neurons recorded in the virtuality task and for neurons recorded in the cued delay task. Neurons recorded at ML position > 900μm were excluded from the analysis as they were considered to be in the substantia nigra pars compacta (SNc).

### Reinforcement learning at multiple timescales

In standard reinforcement learning, the value of a state *s* under a given policy *π* is defined as the expected sum of discounted future rewards:

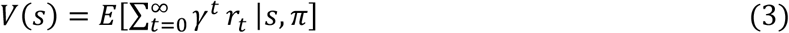

The discount factor *γ* (whose value is between 0 and 1) is a fixed factor at each time step devaluating future rewards. This exponentially functional form for the temporal discount is not arbitrary. This temporal discount is naturally produced by the TD learning rule, a bootstrapping mechanism that updates the value estimates using the experienced transition from *s* to *s*^′^ with reward *r* :

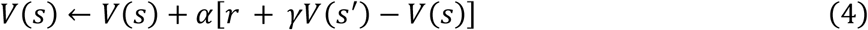

where *α* is the learning rate. This update process converges to the values defined above under very general conditions ^19^ and has been experimentally proven to be an extremely robust and efficient learning rule for Deep RL systems ^22,84^.

After convergence, the value *V*(*s*) can be rewritten by taking the sum and the discount factor outside of the expectation:

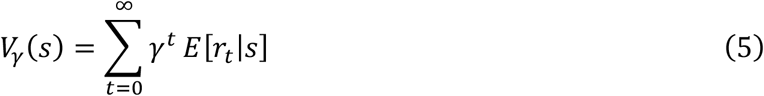

Where we have added a *γ* subscript to the value to indicate that the value is computed for that particular discount, and we have omitted the dependence of the expectation on *π* for simplicity. This last expression reveals a very useful property: *V*_*γ*_(*s*), as a function of the discount *γ* ∈ (0,1), is the unilateral Z-transform of *E*[*r*_*t*_|*s*] as a function of future time *t* ∈ (0, ∞), with real-valued parameter *γ*^−1^ (i.e., the discrete-time equivalent of the Laplace transform^85^). Since the Z-transform is invertible, in the limit of computing values with an infinite amount of *γ*’s, the agent can recover the expected rewards at all future times 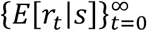 from the set of learned values {*V* _*γ*_ (*s*)}_*γ*∈(0,1_) :

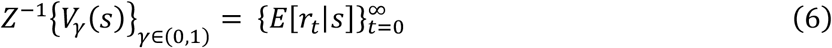

Thus, if the agent performs TD learning with an infinite amount of discounts, the converging points of the TD backups would encode not only the expected sum of discounted rewards, as in traditional RL, but also the *expected reward at all future timesteps*, though the latter lies in a different space, analogous to the frequency and temporal spaces of the Fourier transform.

### Decoding Tasks

The three tasks in Fig. 2c-e were designed with a similar structure. In the three tasks, the policy gradient (PG) network is composed of 2 Fully Connected layers of 32 units each, separated by ReLU nonlinearities. The PG network receives in its input the values learned by TD-learning and reports in its output the corresponding estimate for each task. Values were learned using tabular TD-learning as indicated in the previous section. In Fig. 2c-e and Extended Data Fig. 1, the PG network was trained across 1,000 episodes. The precise structure of each episode depends on the task (see details below). In general, in each episode the agent learns values from scratch using TD-learning for a specific experimental condition (i.e. a Markov decision process, or MDP), and the PG network maximizes its reporting performance across episodes. Thus, for each episode *i*, the policy (*π*_*θ*_) is a map from the learned multi-timescale values 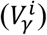 to actions (*a*_*i*_). The parameters (*θ*) of the PG network are optimized to maximize reporting accuracy across episodes (the specific measure to report depends on the experimental condition). The parameters were learned by optimizing the traditional policy gradient loss, using an Adam optimizer with a learning rate of 0.001 to maximize the task-specific expected return *J*(*π*_*θ*_) of the policy *π*_*θ*_:

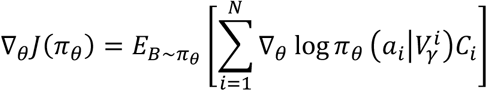

where *B* is a batch of *N=100* episodes and C_*i*_ is a reinforcement learning binary signal indicating whether the report (*a*_*i*_, the output of the network) was correct or incorrect for episode *i*, given the learned multi-timescale values 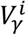 . To tackle the exploration-exploitation problem we extend the policy using *ϵ*-greedy, with *ϵ* = 0.3 (performance is reported with *ϵ* = 0).

In Task 1 (Fig. 2c, Extended Data Fig. 1a-c), in each episode a discrete reward time *t*_*R*_ is sampled between 1 and 15 and a discrete reward magnitude R sampled between 1 and 15. This defines a Markov Decision Process (MDP) shown in Extended Data Fig. 1a. For this MDP, TD-learning was used to learn the value of the first state of the MDP *s*, which we will refer to as the “cue”. In all tasks, the value of the cue was learned using one, two or three discount factors (γ) from the set {0.6,0.9,0.99}, depending on the experimental condition. The results indicated as ‘Three γ’ corresponds to the discount factors [0.6,0.9,0.99]. Since there is noise in the simulation (see below), the results indicated as ‘One γ’ corresponds to the top performer over three identical discount factors ([0.6,0.6,0.6], [0.9,0.9,0.9], [0.99,0.99,0.99]) and analogously for the results indicated as ‘Two γ’. After performing TD-learning, the values are fed as input into the PG network whose output is the guessed reward time (the network has 15 discrete actions, corresponding to reporting reward times from 1 to 15). Performance was evaluated as the fraction of correct responses across test episodes (1 for estimating the correct reward time, 0 otherwise). We show the performance of the PG network as it is trained in Extended Data Fig. 1c. In Extended Data Fig. 1j-l we show a similar experiment but using two reward times and reward magnitudes in the MDP.

In Task 2 (Fig. 2d, Extended Data Fig. 1d-f), the structure of each episode was as in Task 1 but with a discrete reward time *t*_*R*_ sampled between 1 and 8 and a discrete reward magnitude *R* sampled between 1 and 4. The learned values were input into a PG network with 32 possible discrete outputs, representing the 32 possible hyperbolic values obtained in all the possible experiment (4 possible reward magnitudes × 8 possible reward times):

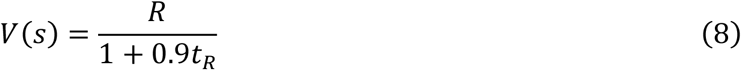

Performance was evaluated as the fraction of correct responses across episodes.

In Task 3 (Fig. 2e, Extended Data Fig. 1g-i) we use the MDP shown in Extended Data Fig. 1g while keeping *R* fixed at 1 but varying *t*_*R*_ and the number of times (*N*) that the full MDP has been experienced by the agents. Since TD-backups are performed online after every transition, *N* is proportional to the total number of TD-backups. The (possibly incomplete) learned values at *s* from these *N* experiences were fed into the PG network (Extended Data Fig. 1h) which was trained across episodes to optimize the reporting performance of *t*_*R*_ .

We also evaluate learning in incomplete-information situations using the MDP shown in Extended Data Fig. 1m-o. In each episode, the length of the two branches is uniformly sampled from 5 to 15 (if they are the same, they are re-sampled until being different). Thus, in each episode, there is a shorter branch and a longer branch. Each branch is experienced a random number of times (*N*) sampled from a uniform distribution with the range of 1 and 99 [denoted by Uniform(1,99)]. Thus, the number of TD backups performed for the two branches could be highly asymmetric. The learned values (with one or multiple discounts) were fed as input into the PG network with a binary output indicating which path was the shortest one, performance was evaluated as the fraction of correct responses (Extended Data Fig. 1o). Single-timescale agents can incorrectly believe that one branch is shorter than the other one if it has been experienced more often, but multi-timescale agents can determine the distance to the reward independently of the asymmetric experience.

In all tasks, the TD-learning process was corrupted by noise. In each episode, the learning rate was sampled from a normal distribution with mean of 0.1 and variance of 0.001 [denoted by *𝒩*(0.1,0.001)] and the number of TD backups was sampled from Uniform(59,99) (except in the tasks with incomplete learning, e.g. Fig. 2e). This variability was included to make sure that the decoder learns robust decoding strategies instead of just memorizing the exact values of each experimental condition. For example, as we argued in the main text, with one discount, the value of a temporally close small reward is similar to the value of a temporally far high reward, so reward time cannot be disentangled from reward magnitude. However, although these two values are similar, they are not identical, so a decoder with enough precision could learn to memorize them in order to report reward time. Introducing a small amount of random noise in the learning process assures robustness in the evaluation of the reporting performance.

### Recovering temporal information before TD learning converges

In Extended Data Fig. 2 we illustrate intuitively why the temporal information is available before TD learning converges for multi-timescale agents (experiment in Fig. 1e). Consider the two experiments in Extended Data Fig. 2a, one with a short wait between the cue and reward (pink) and one with a longer wait (cyan). For a single timescale agent (Extended Data Fig. 2b), the value of the cue depends not only on the experiment length but also on the number of times that each experiment has been experienced (*N*, the number of TD-backups). Thus, for a given set of learning parameters (learning rate, discount factor, timestep length and reward magnitude), the single-timescale agent can incorrectly believe that the cyan cue indicates the shorter trajectory, if it has been experienced more often (left part of the plot). However, as we show theoretically in this section, since temporal information is encoded *across* discount factors for a multi-timescale agent, multi-timescale agents can determine reward timing independently of *N*. In Extended Data Fig. 2c, the patterns of three dots highlighted with rectangles are indicative of the reward time and are only affected by the learning parameters by a multiplicative factor. Indeed, when we plot the multi-timescale values as a function of the number of times that the experiments are experienced (*N*, Extended Data Fig. 2d-e), we see that the pattern across discounts is maintained, enabling a downstream system to robustly decode reward timing.

The following is a theoretical proof of this advantage. Consider a multi-timescale agent performing TD learning on the trajectory *s* → ⋯ → *s*_*T*_ in which there is no variability in outcome timing (i.e., non-zero outcomes always happen at the same states, but their magnitude can be stochastic) and all rewards are positive. Under these assumptions, the agent is able to decode reward timing if it has access to 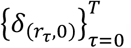, the future times at which outcomes *r*_*τ*_ are non-zero given the current state, where 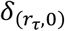 is a Kronecker delta function that is equal to 1 if *r*_*τ*_ is zero and equal to 0 otherwise. *At any time during TD learning*, the value estimate for *s* computed with TD learning can be written with the following general expression (note the absence of the expectation):

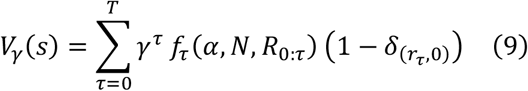

where *f*_*τ*_ (*α, N, R*_0:*τ*_) is a non-zero scalar that depends on *τ*, on the learning rate *α*, on the number of times the trajectory has been experienced *N* and on the history of outcome magnitudes experienced in the past *R*_0:*τ*_. This decoupling shares similarity with the successor representation ^62,86,87^. Crucially, *f*_*τ*_(*α, N, R*_0:*τ*_) does not depend on *γ*, so, at all times during learning, it holds that:

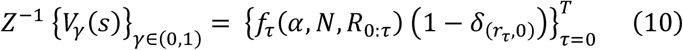

Since *f*_*τ*_(*α, N, R*_0:*τ*_) is non-zero for all *τ*’s and 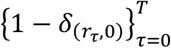 is only non-zero at *τ*’s in which a reward happens, the non-zero values of the right-hand side expression indicates the future reward timings. In other words, applying the inverse transform *at any time during learning* to the multi-timescale estimate {*V* _*γ*_ (*s*) _*γ*∈(0,1)_} gives an expression whose non-zero values are the future outcome timings. In summary, in the absence of timing stochasticity the multi-timescale agent can recover future outcome timing before TD converges, a capability that is not present in single-timescale agents.

### Myopic learning bias: branching task

In Fig. 2f, we present a simple MDP to highlight the myopic learning bias during training. In each episode, the agent learns from 3 trajectories: one that moves up at *s*, another that moves down at *s* but up at state *s’*, and one that moves up at *s* and *s’*. Since rewards are stochastic, the information that the agent gets on each episode is incomplete. To evaluate how well the agent acts given limited information, we average performance over the following procedure: (1) sample rewards along the three trajectories mentioned before, (2) learn the Q-values (until convergence) for *s* and *s’* using the rewards from the sampled trajectories and (3) choose the actions that maximize the Q-values. Performance is then measured as the proportions of right decisions across 10,000 iterations of this procedure. In Fig. 2f we evaluate performance as the fraction of episodes in which the Q-value of the branch with the deterministic reward is higher than the Q-value of the branch without the deterministic rewards.

To evaluate the multi-timescale agent of Fig. 2b on this task, we followed a similar procedure. In each episode, we randomize the identity of the top and bottom branches after the bifurcation, which defines an episode-specific MDP. For each episode-specific MDP, the agent performs Q-learning until near convergence using the 3 trajectories mentioned in the previous paragraph. The Q-values at the current state (*s* or *s’*) are fed into the policy learning architecture of Fig. 2b, which outputs the decision to move up or down in the episode-specific MDP. The policy-learning network is trained across episodes to produce actions that maximize overall task performance. For the single-discount agent, we report the maximum performance over the agents with discounts [0.6,0.6] and [0.99,0.99], which achieve a performance of 77±2% and 83±1% respectively. For the multi-discount agent, we use the discounts [0.6,0.99], which achieves a performance of 94±1%. The error bars correspond to the s.e.m. across 500 episodes in a validation set.

### Myopic learning bias: navigation task

Previous theoretical work showed that a myopic discount in RL can serve as a regularizer when approximating the value function from a limited number of trajectories ^88^. In Extended Data Fig. 3 we highlight the fact that the benefit of the myopic discount is contingent upon the distance between the current state and significant environmental events. Consider the simple navigation scenario depicted in Extended Data Fig. 3a. The agent’s motion is random and isotropic, garnering a minor random reward from a normal distribution with mean 0 and s.d. 0.01 in each step and three more substantial rewards upon reaching the areas denoted by fire (*r* = –4) and water (*r* = 2) symbols. We evaluate how well the agent can determine the true value function (under a discount factor *γ* = 0.99) under the aforementioned stochastic policy. Crucially, the agent must perform this task after experiencing only a limited number of trajectories. The grey arrows show an example trajectory, with the actual and estimated values for these trajectories shown in Extended Data Fig. 3b.

We evaluate accuracy using the Kendall rank correlation coefficient between the true value function in the entire maze and the value estimates. The Kendall coefficient measures the fraction of concordant pairs between the two value functions (across all pairs of states in the maze). For every pair of states, it computes whether the two value functions agree on which element of the pair is the larger one. Note that this measure of accuracy is behaviorally more relevant than alternative accuracy measures that compare the absolute magnitude of values across states. In other words, for an agent navigating the maze, it is more important to be accurate on the relative values of alternative goal states than on their absolute values. Consider the trajectory shown in Extended Data Fig. 3b. For this trajectory, the myopic estimate (using a discount factor *γ* = 0.6, green) clearly provides a better estimate of the true value function (grey) than using the true discount factor *γ* = 0.99 (brown). We can quantify that the myopic estimate is a better approximation of the true value function by evaluating the agreement between pairs of states along the estimated and true curves (i.e. by computing the Kendall coefficient).

In Extended Data Fig. 3c-d the agent learns from N randomly sampled trajectories starting either in the lower half (blue) or upper half (red) of the maze. The values for the states in the *N* sampled trajectories are learned until convergence using the rewards and transitions in the sampled trajectories. After convergence, we compute the Kendall rank correlation between the estimates and the true value function, and report performance as the average correlation across 10,000 sets of *N* sampled trajectories. Extended Data Fig. 3c shows that when learning from two randomly sampled trajectories, the estimates of the value function using a myopic discount factor are more accurate than far-sighted discounts when crucial events are in the near future (i.e the trajectories start in the lower half of the maze, blue curve in Extended Data Fig. 3c). This result agrees with the intuition built in Extended Data Fig. 3c when learning from a single trajectory. However, if the agent is distant from important events (i.e. trajectories starting in the upper half of the maze, red curve), the myopic estimates approach the noise level, while estimates with larger discount factors are more accurate. As expected, with the accumulation of more data from the environment, that is, more trajectories, the far-sighted estimate progressively aligns with the true value compute with *γ* = 0.99 in the entire maze (Extended Data Fig. 3d)

### Myopic learning bias: networks with discount factors as auxiliary tasks

An alternative way to leverage multi-timescale learning benefits, in contrast to the architecture presented in Fig. 2b, is to employ them as auxiliary tasks (Fig. 2g, top). These networks only act according to the value of a single behavioral timescale, but concurrently learn about multiple other timescales as auxiliary tasks to enhance the representation in the hidden layers, which allows them to obtain superior performance in complex RL environments ^60,89^. This approach is similar to Distributional RL networks that learn the quantiles of the value distribution but act according to the expectation of that distribution ^41^. Notably, we show that the auxiliary learning timescales display the myopic learning bias highlighted so far. In the Lunar-Lander task (Fig. 2g, bottom) where the agent must land a spacecraft, Q-values computed using a myopic discount provide a more accurate representation of the future when the agent is close to the landing site (blue), whereas the opposite holds when the agent is far from the landing site (red).

In the Lunar Lander environment in Fig. 2g, the state space consists of eight elements, including the position and velocity of the lander, its angular position and angular velocity, as well as an additional input related to the contact with the ground. The action space is composed of four actions: doing nothing and activating one of three different engines. The agent is a Deep-Q-network ^3^ (DQN) with two hidden layers of 512 units each, separated by ReLU activation functions. In addition to the Q-values that control the agent, the network has Q-values for 25 additional discounts factors equally spaced between 0.6 and 0.99. Thus, if there are |a| actions in the environment, for each discount the network has |a| additional output units. All sets of |a| units (one for each discount) use the Huber (i.e. Smooth L1, *β=1*) Q-learning loss function with its corresponding discount. All the auxiliary Q-learning losses update the action that was actually chosen in the environment by the behavioral units, and thus all of them learn the consequences of the behavioral policy, but using different discount factors. The total loss function uset to train the network averages the Q-learning losses of all the discount factors. To train the DQN, we use a learning buffer of 20,000 samples, a learning rate of 10^-3^ and a batch size of 32. As in traditional DQNs, we use a target network to compute the TD target, which is updated every 1,000 samples with the weights from the policy network to stabilize the learning process. For exploration, the agent uses a linearly decreasing ε-greedy policy that goes from *ε = 1*.*0* at the first sample to a minimum value of *ε = 0*.*01* after 40,000 samples.

Our goal is to compute the degree to which Q-values computed with alternative discounts can capture the true Q-value of the behavioral policy. The multi-timescale DQN uses a behavioral discount *γ*_*beh*_ = 0.99, and its policy is produced by choosing actions that maximize the Q-values with that discount factor. As in the navigation scenario presented in the previous section, our hypothesis is that, when important events lie in the proximal future (here, close to the landing site), the Q-values learned using myopic discounts capture the true behavioral Q-value more accurately, while far-sighted discounts are more accurate when important events lie in the distant future (far from the landing site).

Under the policy of the DQN (*π*_*DQN*_), the true value of state *s* is:

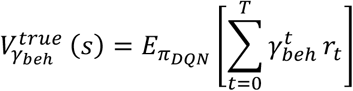

If the DQN has perfectly learned the Q-value of state *s*, then the estimate *Q*_*γ*_(*s, a*_*beh*_) of the DQN should be equal to 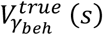, where *a* _*beh*_ is the action produced by the DQN at *s*. We evaluate accuracy as the degree to which the estimated *Q*_*γ*_(*s, a*_*beh*_) captures the true 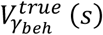, and compare accuracy across the auxiliary discount factors.

After training the network for 50,000 samples (and achieving close-to-optimal performance), we compute 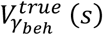 empirically across states by recording the actual discounted sum of rewards obtained by the agent when departing from state *s*. We calculate 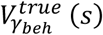 empirically for 25,000 states. Then, we compare, across states, the empirically calculated 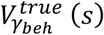 with the Q-values produced by the DQN at those states.

To measure Accuracy, we use the Kendall rank correlation as in the previous section. The Kendall correlation measures the fraction of concordant pairs between samples from 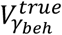 and from the estimated *Q*_*γ*_, across pairs of states. As in the navigation scenario presented in the previous section, for an agent deciding which state to navigate to, it is more important to be accurate on the relative values between pairs of states than on the absolute value of individual states. Therefore, the Kendall correlation is behaviorally more relevant than other accuracy metrics that compare the absolute magnitude of 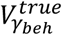 and *Q* _*γ*_ (e.g. 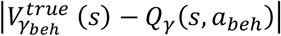).

Given that the environment and the training process are stochastic, we report the accuracy by averaging over 10 randomly initialized networks.

### Cued delay task

All the data in the experiments with mice were collected in the previous study ^68^. The experimental details including the surgical procedures, behavioral setup, and the behavioral tasks have been described there ^68^. We will here focus on the task description as our analysis includes task conditions that were not analyzed in the previous study.

Mice were head-fixed on a wheel in front of three computer monitors and an odor port. At trial onset, the screens flashed green to indicate the beginning of the trial. After *t* = 1.25s, an odor cue was delivered. This reward delay cue was one of four possible odors, and each cue was associated with a unique reward delay chosen from 0.6, 1.5, 3.75 or 9.375 seconds. The association between odor and reward delay was randomized across mice. The inter-trial interval was adjusted depending on the reward delays such that the trial start cues were spaced by 17-20s. Mice performed 81.4 ± 12.5 trials (mean ± s.d.) per session across the 36 sessions in which neurons were recorded in the task.

### Approach-to-target virtual reality (VR) task

We refer the reader to the prior study for details on the experimental procedures ^68^. Mice were also trained in additional conditions, which we do not analyze in the present study, including teleport and speed modulation in the virtual reality scene.

Here, we analyzed single neuron recordings in the sessions with no teleport or speed manipulation and in the open-loop condition. Mice were free to locomote, but their motion did not affect the dynamics of the visual scene. After scene motion onset, the visual scene progressed at constant speed until the reward was delivered after 7.35s.

Mice performed 58.8 ± 21.7 trials (mean ± s.d.) per session across the 60 sessions in which neurons were recorded in the task. Spiking activity was convolved with a box filter of length 10 ms. When plotting neural activity, we further convolved the responses by a causal exponential filter (*e*^*-0*.*05dt*^). Spiking rate traces across neurons were normalized using a modified z-score. The mean was taken as the average firing activity cross the first 1.5s and the standard deviation across the entire 4.35s.

### Fitting neural activity in the cued delay task

For the cued delay task, we fit the responses of single neurons to the delay cue (calculated as the firing rate in the time interval 0.1*s* < *t* < 0.4*s* after the cue onset, see shaded area in Fig. 3c) using two discounting models as in ref[^63^], the classic exponential model and a hyperbolic model. For the exponential model, we fit the responses to a cue predicting a reward in *τ* seconds by:

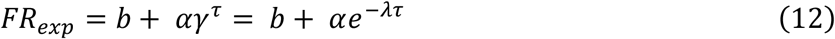

The discount factor *γ* can also be expressed as a discount rate λ and vice versa: *λ* = − ln *γ* or *γ* = *e*^−*λ*^. The discount factors fitted to data are always expressed in units of *seconds*, that is the discount factor is the devaluation one second into the future.

For the hyperbolic model we used a standard model for hyperbolic discounting in which the parameter *k* controls discounting:

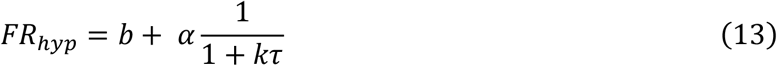

We fitted both models by minimizing mean squared error (the *fit* function in MATLAB). For both models we constrained the baseline and gain parameters such that 0 < *b* < 40 and 0 < *α* < 40. For the exponential model, the discount rate was constrained such that 0.0001 < *λ* < 20 and for the hyperbolic model, the discount parameter was constrained such that 0 < *k* < 20. Note that all the parameters are fitted independently for each single neuron.

To characterize the robustness and significance of our estimated parameters we used a bootstrap procedure. For each run, we split the trials in half and fit the models independently on each half. We computed for each split the explained variance using the other half of the data (Extended Data Fig. 4c-d) and correlated the inferred parameter values for each neuron across both splits (Extended Data Fig. 4f-g).

We restricted our subsequent analysis to neurons that had a positive explained variance on the test set (*n=17* neurons excluded), an average firing rate in the cue period over the 4 delays above 2 spikes/s (*n=11* neurons excluded) and with medio-lateral distance above 900μm (*n=4* neurons excluded). Non-selected neurons are shown in Extended Data Fig. 4b. Poorly fit neurons often were non-canonical dopamine neurons who also did not exhibit a strong reward response.

### Decoding expected reward timing from population responses

The vectorized prediction error allows us to directly decode the expected timing of reward given the cue responses^45^. The value at time t is given by:

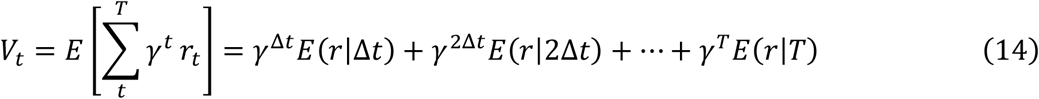

In the cued delay task, at the time of the cue indicating reward delay, the response of dopaminergic neurons is driven by the discounted future reward. The reward prediction error *δ*_*t*_ = *r*_*t*_ + *γ*^Δ*t*^*V*_*t*+1_ − *V*_*t*_ becomes simply *δ*_*t*_ = *γ*^Δ*t*^*V*_*t*+1_ + *cst* as there is no reward delivered at the time of the cue 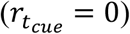 and the reward expectation before the reward cue delivery is identical across conditions (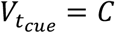; where *C* is a constant). Thus, the TD error at the time of reward delay cue 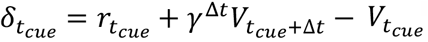 becomes 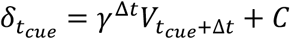 and if we assume the constant is 0 or the TD-error is baseline subtracted, at convergence the prediction error is given by:

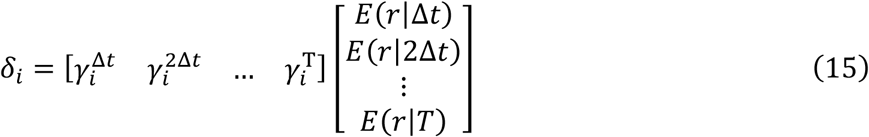

In single timescale RL, the temporal information is collapsed, and it is not possible for the system receiving the learning signal (the striatum in this case) to untangle the signal. However, in a distributed system learning at multiple timescales the reward expectation *E*(*r*|*t*) is encoded with multiple discount factors *γ*_*i*_ :

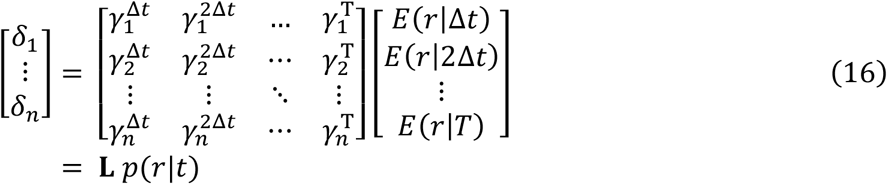

The temporal information about reward timing is now distributed across neurons and if the tuning of individual neurons is sufficiently diverse, we can write:

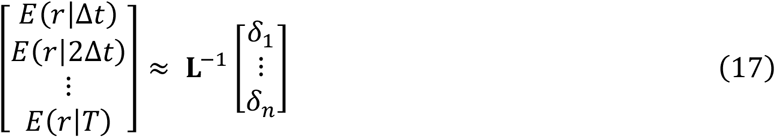

Where **L**^−1^ is the approximate pseudo-inverse of **L** such that **L**^−1^**L** ≈ *I*. In practice, the matrix **L** is not very well conditioned as the rows of the matrix are exponentially decaying functions, so the right side (further in the future) is sparsely populated (see Fig. 3i and Extended Data Fig. 5a-c). We therefore will need to use a regularized pseudoinverse.

To invert the discount matrix **L**, we use the regularized Singular value decomposition (SVD) approach similar to the one proposed in ref[^45^]. We then normalize the resulting prediction in order to constrain it to be a probability distribution (*p*(*r*|*t*) > 0, *for all t* and **∑**_*t*_ *p*(*r*|*t*) = 1). More specifically, the regularized SVD approach corresponds to optimizing:

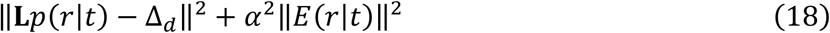

The standard SVD of the discount matrix can be written as:

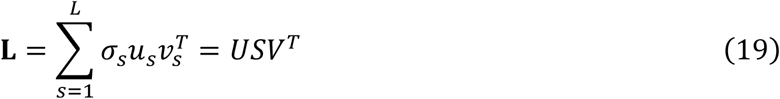

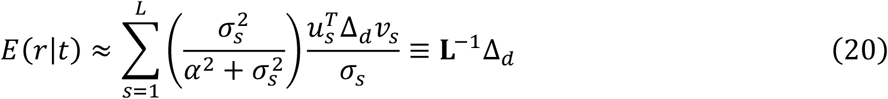

where Δ_*d*_ = [*δ*_1_ … *δ*_*N*_]^*T*^. The smooth regularization introduced by the Tikhonov regularization through the parameter *α* (which we can choose by inspection of the distribution of singular values *σ*_*s*_, see below) is more robust than a strict truncated SVD in which we only take a number of factors and set the remaining ones to zero. An alternative approximation to this inverse problem is Post’s approximation ^57,62^. It relies on evaluating higher order derivatives and lacks robustness if the Laplace space is not sampled with enough precision (i.e. not enough neurons tiling the γ space).

The procedure in the previous section allows us to estimate the discount factor independently for each neuron. We then choose a discretization step Δ*t* = 100*ms* and a temporal horizon *T* = 12*s* over which to make the prediction. This allows us to construct the discount matrix **L** shown in Fig. 3h for the exponential model and Extended Data Fig. 6c for the hyperbolic model. In order to choose a suitable value for the regularization parameter *α* we perform the regular SVD on the discount matrix **L** and assess the values at which the singular values become negligible. We choose a value of *α* that corresponds to the transition between large singular values and negligible ones (see Extended Data Fig. 5b). Using this approach, we used *α* = 2 in our decoding analysis.

For each delay, we construct a pseudo-population response Δ_*d*_ across the recorded neurons. For each bootstrap, we take the mean activity for each cue, subtract the inferred baseline parameter *b*, and normalize the maximum response to 1. To assess the robustness of the predictions, we use the mean responses and baseline from half the trials to construct Δ_*d*_ and use the estimated discount factors from the other half of the trials to estimate **L**^−1^ and we repeat this approach for each bootstrap (*n*_*predictions*_ = 200). In the figures (Fig. 3k and Extended Data Fig. 6d,f and 9c), the thin lines correspond to the predictions from individual bootstraps and the thicker line to the average of these predictions. For shuffle control, we randomize the identity of the neurons in the pseudo-population response Δ_*d*_. This means that in the shuffle control a given neuron is not decoded with its corresponding weights but by a random row of the decoding matrix **L**^−1^.

In order to ensure that the prediction corresponds to a probability distribution, we normalize the resulting prediction of reward timing. We first set the probability of obtaining a reward to zero for all times in which the prediction was negative, then we normalize the distribution to be a valid probability distribution (such that the probability mass over *t* ∈ [0,12] sums to 1).

For the time decoding using a single average discount factor, we use a different approach. The inversion procedure would not work as the discount matrix would be of rank 1. Instead, if we assume a fixed known reward size and a single discount factor, the response of individual neurons would correspond to different estimates of the reward timing. For each bootstrap we can estimate the expected reward timing for each neuron. For a given firing rate FR for the held out data, we can estimate the reward timing using the parameter estimates from the trained data. The baseline *b*_*i*_ and gain *α*_*i*_ parameters are specific to each neuron while the discount factor *γ* is the average discount factor across all the neurons. The expected reward timing for neuron *i* is given by the following equation:

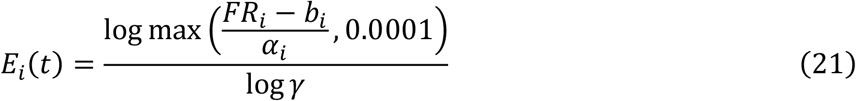

Together, the neurons provide a distribution of expected reward timing with each neuron predicting a sample of the distribution of expected reward times. The average distribution is obtained by averaging the distributions across all the bootstraps, excluding predicted reward times beyond 12 seconds and normalizing the distribution to be a probability distribution. Similarly to the SVD-based decoding, in Extended Data Fig. 5f the thin lines correspond to the predictions from individual bootstraps and the thicker line to the average of these predictions.

### Quantifying reward timing decoding accuracy

In order to quantify the reward timing decoding accuracy, we used the 1-Wasserstein distance (or earth mover’s distance) between distributions as our metric. We used the 1-Wasserstein distance as the difference in support between the predicted reward timing distribution (probability mass as most locations) and the single true reward timing (probability mass at a single location) is not conducive to using the KL-divergence.

For each bootstrap, we generated *n* = 100,000 samples from the predicted reward timing distributions and computed the 1-Wassertsein distance between the predicted reward timing and the true corresponding reward delay (using the MATLAB function *ws_distance* from https://github.com/nklb/wasserstein-distance). For each condition (exponential fit, hyperbolic fit, average discount factor, simulation fit and their associated shuffled predictions) we obtained a distribution of 1-Wasserstein distances across the bootstraps (*n* = 200). To assess the significance of the differences in reward timing predictions across conditions, we used the one-tailed Wilcoxon’s signed rank test (using the MATLAB function *signrank*).

### Fitting neural activity in the VR task

To quantify the heterogeneity of discount factors in the VR task, we fit the neural activity in the last 4.30 seconds (*t* = 3.05 seconds after scene motion onset) of the approach to reward period in which the ramping activity was most pronounced. In order to assess the robustness of the fit, we used a bootstrap procedure in which for each bootstrap (*n*_*bootstrap*_ = 100), we partition the trials in two halves and compute the two average PSTHs using *dt* = 0.1 second as our discretization step. We then compute the mean value of the parameters across all bootstraps. We limit our analysis to neurons whose firing rate over the analysis period is larger than 2 spikes/s. We fit the two models (common value function and common reward timing expectation) to this data.

In the VR task, the expectations vary smoothly as a function of time and distance and we therefore use the discretized formulation of the TD error for continuous time in our fits ^69,71^:

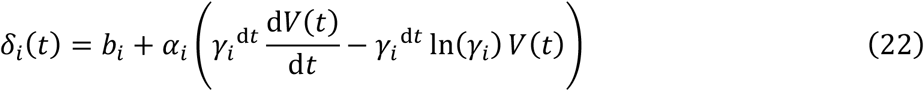

Although this formulation is also discretized as the standard formulation of the TD error, the presence of the derivative 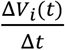 (which is computed numerically) improves the stability of the fitting procedure. The two models differ in whether value function is estimated directly (and shared across neurons) or indirectly (and distinct across neurons). The discount factor is also in units of *seconds*, allowing for a comparison with the values estimated in the cued delay task.

### Common value function model

In the common value function model, *V(t)* is common across neurons and is directly fitted by the optimization procedure which minimizes:

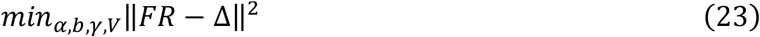

With,

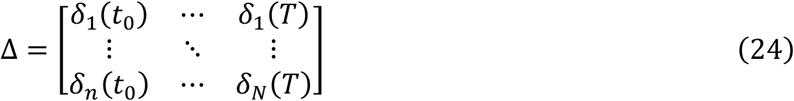

We fit the gains, baseline, and discount factors of individual neurons (*α*_*i*_, *b*_*i*_ and *γ*_*i*_ respectively) and the join value function *V* using a constrained optimization procedure (*fmincon* in MATLAB, *α*_*i*_ ∈ [0.05,50], *b*_*i*_ ∈ [0.05,12], *γ*_*i*_ ∈ [0.05,0.999999], and *V* ∈ [0.05,5]).

### Common Reward Expectation model

In the common reward expectation model, the reduction in uncertainty in reward timing due to sensory feedback as the mice approach the reward leads to an upwards ramp in the average TD error signal across dopaminergic neurons ^68–70^. In a task like the cued delay task shown in Fig. 3, once the cue has been presented, the time estimation until the reward is based on the internal clock of the mice that suffers from scalar timing (i.e., the standard deviation of the noise in the estimation grows linearly with the estimation time ^49^. In the VR task, there is visual feedback and as the mice approach the reward, the uncertainty is instead reduced (Extended Data Fig. 8a). We also show that this alternative model also provides a similar explanation of ramping diversity as originating from a heterogeneity of discount factors (Extended Data Fig. 8).

We use a joint fitting procedure in which we simultaneously fit the discount factors across neurons and the expected timing of reward as a function of position in the virtual track. Similarly to ^69^, we interpret the ramping in single neurons as originating from the reduction in uncertainty due to the visual feedback as the mice approach the reward. Although each neuron has a distinct discount factor and its own value function, the world model which parametrizes the changes in reward expectation with visual feedback is shared across dopaminergic neurons. This arises as this shared model is the product of the integration of the diverse dopamine signals as well as other neural computations controlling reward expectations ^29^.

Individual neurons therefore act as independent agents estimating value given a shared expectation of reward timing. Each neuron has a distinct discount factor *γ*_i_ with which it computes value given the expected reward timing. We assume that inference has converged and therefore we have the value *V*_*i*_ associated with neuron *i*:

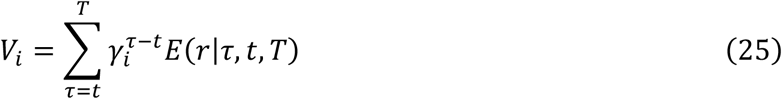

Here, we assume that *E*(*r*|*τ, t, T*) takes the form a folded normal distribution with parameters *μ* = *T* − *t* and (fitted) standard deviation *σ*. The folded normal distribution reflects the weight of the negative component of a normal distribution back onto positive values ^90^. The folded normal distribution formulation leads to the following distribution for the expected reward timing for *τ* > 0 :

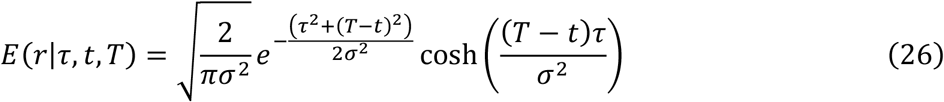

In our analysis, the mean, *μ* = *T* − *t*, is given by the current position in the VR track and the only fitted parameter is the standard deviation *σ*. At each time step we fit a different value of the standard deviation. As observed through the fitting procedure, the standard deviation is initially high and reduces as the mice approach the reward location. This is an indication that similarly than proposed in ^69^ the ramping in the dopaminergic neuron’s activity arises from the reduction in uncertainty due to the visual feedback as the mice approach the reward. We use a slightly different formulation than in ref [^69^] as we require additional flexibility to fit data and specifically need to go beyond the assumptions of Gaussian state uncertainty. Note also that we assume here that the uncertainty is in the timing of the reward rather than in the state.

In order to normalize the contributions of the different neurons, we used a normalized firing rate and therefore only fit the discount factor *γ* and standard deviation *σ* of the reward expectation.

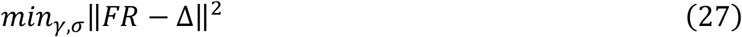

With,

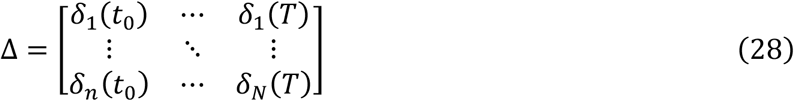

We performed the constrained optimization with the *MATLAB* function *fmincon* and constrain the parameters such that *γ* ∈ [0.001, 0.99] and *σ* ∈ [0.1, 12].

### Comparing parameters across tasks

We used two methods to assess the relationship between the inferred discount factors in the approach-to-reward VR task and the cued delay task. First, we used the mean parameters across bootstraps and computed the Spearman correlation. Next, we computed, for *n* = 10,000 randomly selected (with replacement) pairs of bootstraps, the Spearman correlation between the parameters across the two tasks and plotted the distribution of these correlation.

For the decoding of reward timing using parameters inferred in the VR task, we also used a bootstrap approach. We computed the discount matrix and the decoding matrix for each bootstrap estimate of the discount factors in the VR task.

### Simulations to assess limits on parameter estimation

To assess the contribution of the limits imposed by the number of trials and the stochasticity in firing rates to the accuracy of the reward timing prediction and the similarity of inferred parameters across tasks, we ran a series of simulations with parameters chosen to match those inferred from the data. For the simulation parameters, we use the mean inferred value for the parameters across all the bootstraps for the respective task.

For the cued delay task, we generated for each neuron *n* = 80 trials (*n* = 20 per delay), comparable to behavioral sessions in the task, simulated cue responses by taking samples from a Poisson distribution with a rate parameter corresponding to the value predicted by the exponential discount model for the corresponding reward delay. We used the same procedure as for analyzing the recorded data by performing *n* = 100 bootstrap and fitting the simulated data on random partitions of the data.

For the VR task, we generated for each neuron *n* = 80 trials, comparable to behavioral sessions in the task, by taking samples from a Poisson distribution with a rate parameter corresponding to the predicted activity given equation 22. We then performed the fitting procedures similarly than for the experimental data.

## Acknowledgements

We thank Dr. Samuel Gershman and Dr. John Mikhael for their contributions to the preceding studies and Dr. Mitsuko Watabe-Uchida for advice on task design. We thank the members of the Uchida and Pouget labs including Adam Lowet and Mark Burrell for discussions and comments. We also thank Wilka Carvalho, Gautam Reddy and Torben Ott for their comments on the manuscript. This work is supported by NIH BRAIN Initiative grants (R01NS226753, U19NS113201) and NIH grant 5R01DC017311 to N.U and by a grant from the Swiss National Science Foundation (315230_197296) to A.P.

## Author contributions

P.M., P.T., A.P. and N.U. conceived the project. P.M., H.R.K, A.N.M and N.U designed the electrophysiology experiments. A.N.M and H.R.K. performed the electrophysiology experiments and curated data. P.T. Performed simulations with artificial agents. P.M. Performed analysis of electrophysiological data. P.M., P.T., A.P. and N.U. wrote the paper with input from H.R.K.

## Data availability

The code used for simulations can be found at https://github.com/pablotano8/multi_timescale_RL. The data code for the electrophysiological experiments and the corresponding analysis code will be uploaded to a public repository upon acceptance.

## Extended data figures and tables

**Extended Data Fig 1.**
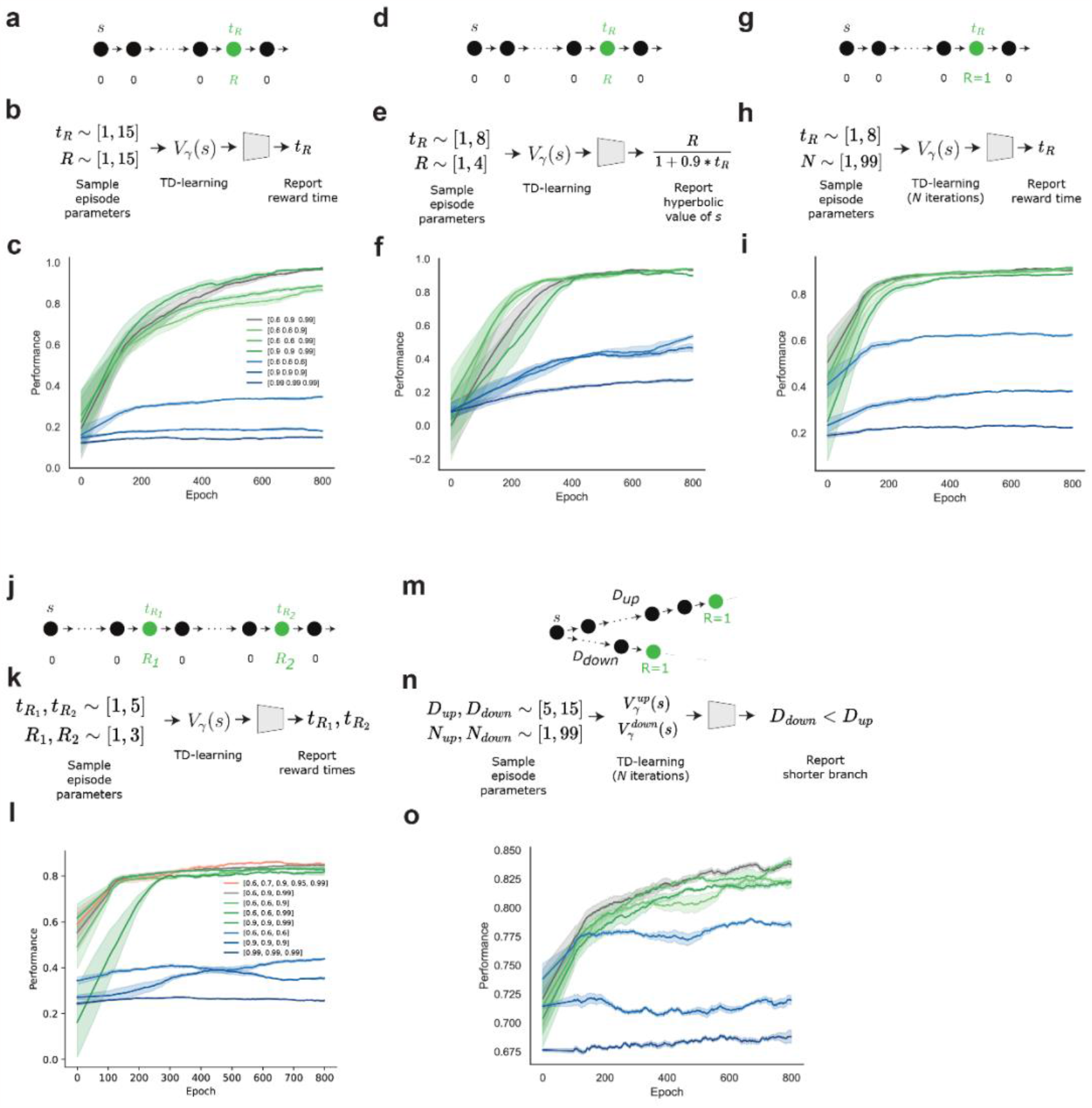
Decoding simulations for multi-timescale vs. single-timescale agents. **(a-c)**. Experiment corresponding to Fig. 2c. (decoding reward timing). **a**, MDP with reward *R* at time *t*_*R*_ . **b**, Diagram of the decoding experiment. In each episode, the reward magnitude and time are randomly sampled from discrete uniform distributions, which defines the MDP in **a**. Values are learned until near convergence using TD-learning. Values with different discount factors are learned independently. The learned values for the cue (*s*) are fed into a non-linear decoder which learns, across MDPs, to report the reward time. **c**, Decoding performance as the decoder is trained. Different colors indicate the discount factors used in TD-learning. **(d-f)**. Experiment corresponding to Fig. 2d. (Decoding value with hyperbolic discount). **d**, MDP with reward *R* at time *t*_*R*_ . **e**, Diagram of the decoding experiment. In each episode, the reward magnitude and time are randomly sampled from discrete uniform distributions, which defines the MDP in **d**. Values are learned until near convergence using TD-learning. Values with different discount factors are learned independently. The learned values for the cue (*s*) are fed into a non-linear decoder which learns, across MDPs, to report the hyperbolic value of the cue. **f**, Decoding performance as the decoder is trained. Different colors indicate the discount factors used in TD-learning. **(g-i)**. Experiment corresponding to Fig. 2e. (decoding reward timing before convergence). **g**, MDP with reward equal to 1 at time *t*_*R*_ . **h**, Diagram of the decoding experiment. In each episode, the reward time and the number of TD iterations (*N*) are sampled from discrete uniform distributions. Values are learned by performing N TD-learning backups on the MDP. Values with different discount factors are learned independently. The learned values for the cue (*s*) are fed into a non-linear decoder which learns, across MDPs, to report the reward time. **i**, Decoding performance as the decoder is trained. Different colors indicate the discount factors used in TD-learning. **(j-l)**. Decoding reward timing in a more complex task. **j**, MDP with two rewards of magnitude *R*1 and *R*2 at times *t*_*R*1_ and *t*_*R*2_. **k**, Diagram of the decoding experiment. In each episode, both reward magnitudes and times are sampled from discrete uniform distributions. The learned values for the cue (*s*) are fed into a non-linear decoder which learns, across MDPs, to report both reward times. **l**, Decoding performance as the decoder is trained. Different colors indicate the discount factors used in TD-learning. **(m-o)**. Decoding length of branches in an MDP during training. **m**, MDP with two possible trajectories. In this example, the upwards trajectory is longer than the downwards trajectory. **n**, Diagram of the decoding experiment. In each episode, the length of the two branches D and the number of times that TD-backups are performed for each branch are randomly sampled from uniform discrete distributions. Then, TD-backups are performed for the two branches the corresponding number of times. After this, they are fed into a decoder which is trained, across episodes, to report the shorter branch. **o**, Decoding performance as the decoder is trained. Different colors indicate the discount factors used in TD-learning. In panels **c, f, i, k** and **o**, the shaded area corresponds to the standard deviation of the estimate over 2 repeats and smoothed of 100 episodes.

**Extended Data Fig 2.**
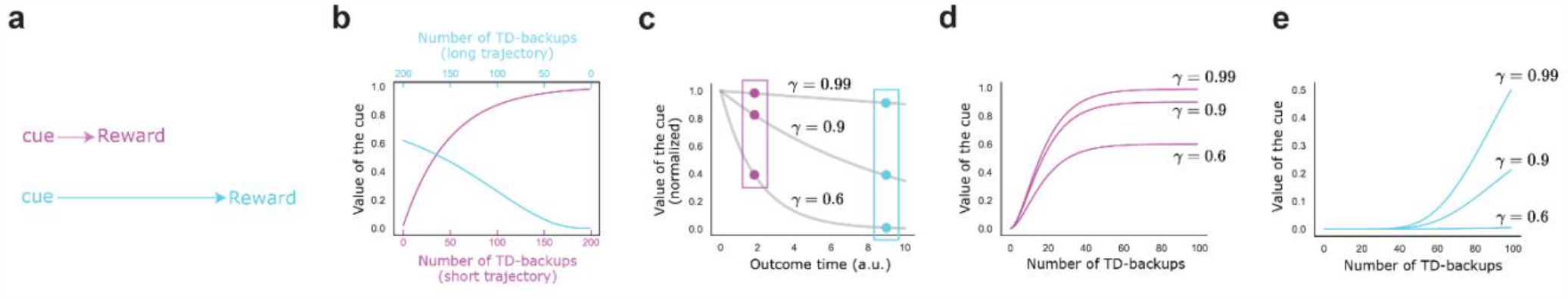
Temporal estimates are available before convergence for multi-timescale agents. **a**, Two experiments, one with a short wait between the cue and reward (pink), and one with a longer wait (cyan). **b**, The identity of the cue with the higher value for a single-timescale agent (here *γ*=0.9) depends on the number of times that the experiments have been experienced. When the longer trajectory has been experienced significantly more often than the short one, the single-timescale agent can incorrectly believe that it has a larger value. **c**, For a multi-timescale agent, the pattern of values learned across discount factors is only affected by a multiplicative factor that depends on the learning rate, the prior values and the asymmetric learning experience. The pattern therefore contains unique information about outcome time. **d**,**e**, When plotted as a function of the number of times that trajectories are experienced, the pattern of values across discount factors is only affected by a multiplicative factor. In other words, for the pink cue, the larger discount factors are closer together than they are to the smaller discount factor, and the opposite for the cyan cue. This pattern is maintained at every point along the x-axis, and therefore is independent of the asymmetric experience, and it enables a downstream system to decode reward timing.

**Extended Data Fig 3.**
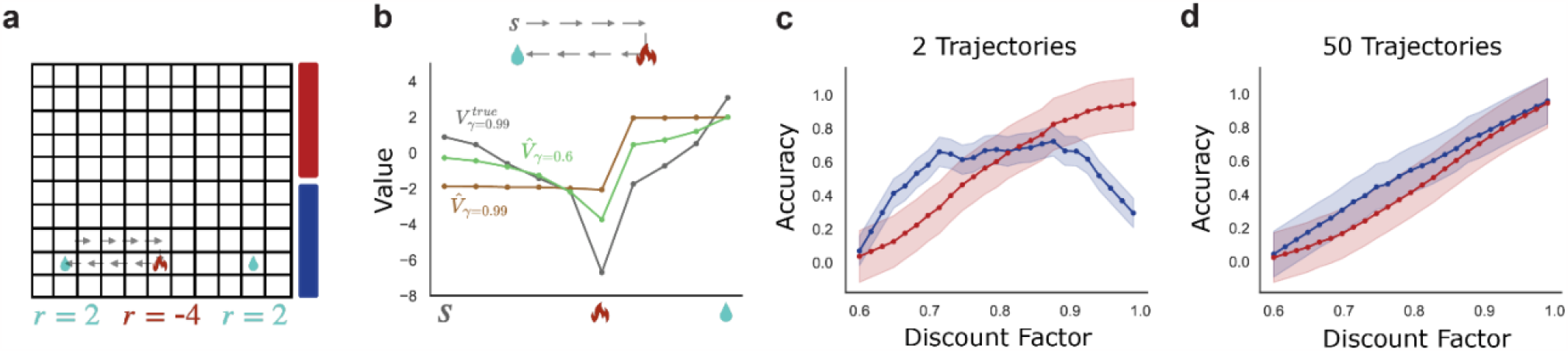
Myopic learning bias. **a**, Maze to highlight the myopic learning bias. Rewards are indicated with water and fire. An example trajectory is shown with transparent arrows. The red and blue bars to the right denote the states in the Lower and Upper half. **b**, True (grey) and estimated (green and brown) values for the example trajectory on top and shown in panel a. In the x-axis we highlight the starting timestep with *s*, the timestep when the fire is reached and the timestep when the water is reached. **c**, Accuracy (y-axis) is measured as the Kendall tau coefficient between the estimate with a specific gamma (x-axis) and the true value function *V*_*γ*_ = 0.99. Error bars are deviations across 300 sets of sampled trajectories. The red (blue) curve shows average accuracy for the states on the upper (lower) half of the maze, indicated with color lines on panel a. **d**, As the sampled number of trajectories increases, the myopic learning bias disappears.

**Extended Data Fig 4.**
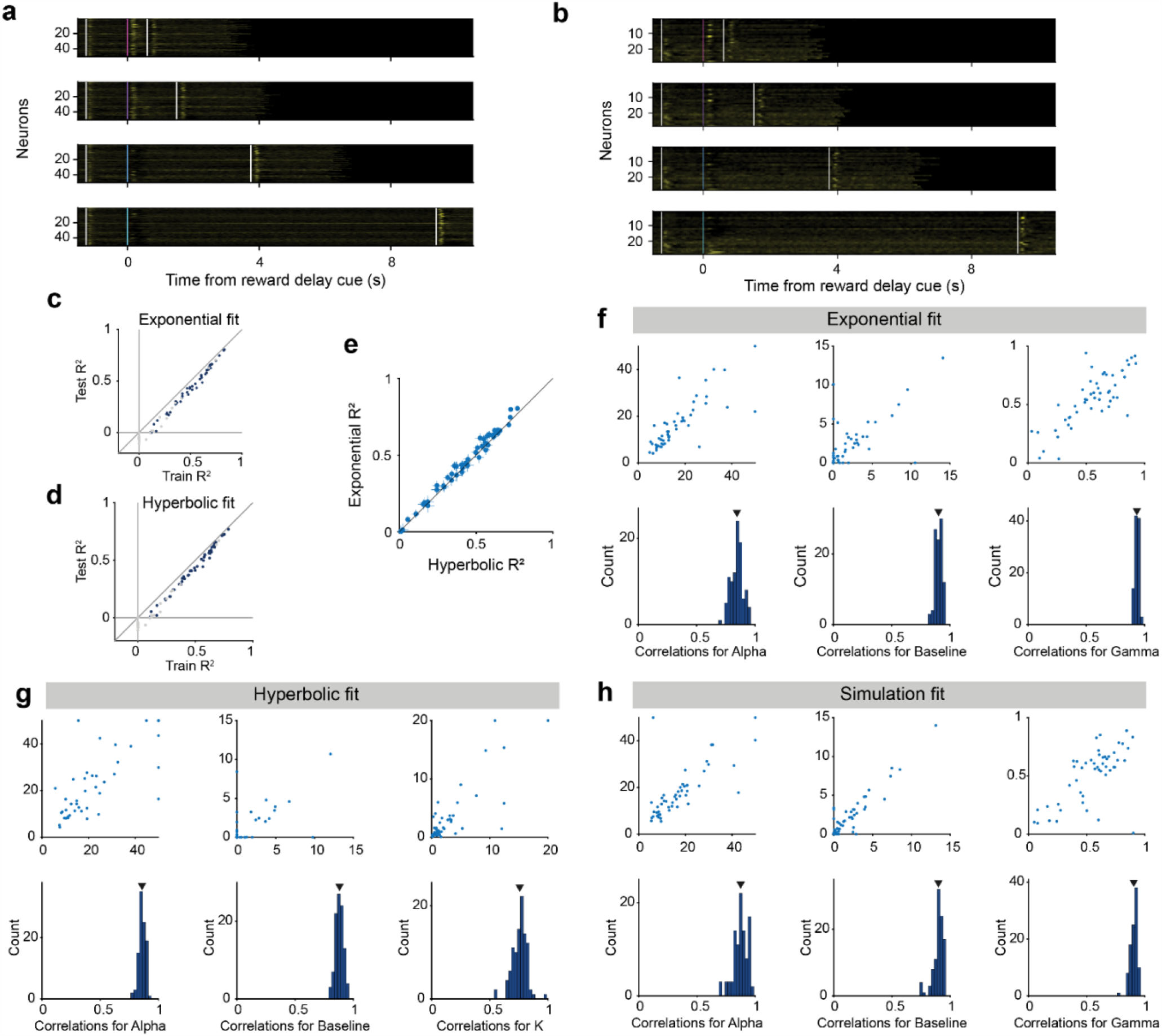
Single neuron responses and robustness of fit in the cued delay task. **a**, PSTHs of single selected neurons (*n* = 50) responses to the cues predicting a reward delay of 0.6s, 1.5s, 3.75s, and 9.375s (from top to bottom). Neurons are sorted by the inferred value of the discount factor *γ*. Neural responses are normalized by z-scoring each neuron across its activity to all 4 conditions. **b**, PSTHs of single non-selected neurons (*n* = 23) responses to the cues predicting a reward delay of (from top to bottom). Neurons are sorted by the inferred value of the discount factor *γ*. Neural responses are normalized by z-scoring each neuron across its activity to all 4 conditions. **c**, Variance explained for training vs testing data for the exponential model. For each bootstrap, the variance explained was computed on both the half of the trials used for fitting (train) and the other half of the trials (test). Neurons (*n* = 13) with a negative variance explained on the test data are excluded from the decoding analysis (grey dots). **d**, Same as panel **c** but for the fits for the hyperbolic model. **e**, Goodness of fit on held-out data for each selected neuron for the exponential and hyperbolic models. The data lies above the diagonal line suggesting a better fit from the exponential model as shown in Fig. 3f. Error bars indicate 95% confidence interval using bootstrap. **f**, The values of the inferred parameters in the exponential model are robust across bootstraps. top row, Inferred value of the parameters across two halves of the trials (single bootstrap) for the gain α, baseline b and discount factor *γ* respectively. Bottom row, Distribution across *n* = 100 bootstraps of the Pearson correlations between the inferred parameter values in the two halves of the trials for the gain α (mean = 0.84, *P* < 1 x 10^-20^), baseline b (v, mean = 0.9, *P* < 1.0 x 10^-32^) and discount factor *γ* (vi, mean = 0.93, *P* < 1.0 x 10^-46^). **g**, Same as panel **e** but for the hyperbolic model with distribution of correlations for the gain α (mean=0.86, p<1e^-26^), baseline b (v, mean = 0.88, P < 1.0 x 10^-28^) and shape parameter k (vi, mean = 0.76, *P* < 1.0 x 10^-11^). **h**, Same as panel **e** and **g** but for the exponential model simulated responses with distribution of correlations for the gain α (mean = 0.86, *P* < 1.0 x 10^-10^), baseline b (v, mean = 0.88, *P* < 1.0 x 10^-24^) and discount factor *γ* (vi, mean = 0.76, *P* < 1.0 x 10^-26^). Note that the distributions of inferred parameters are in a similar range than the fits to the data suggesting that trial numbers constrain the accuracy of parameter estimation. Significance is the highest *p*-value for all the bootstraps for a given parameters assessed via *t*-test.

**Extended Data Fig 5.**
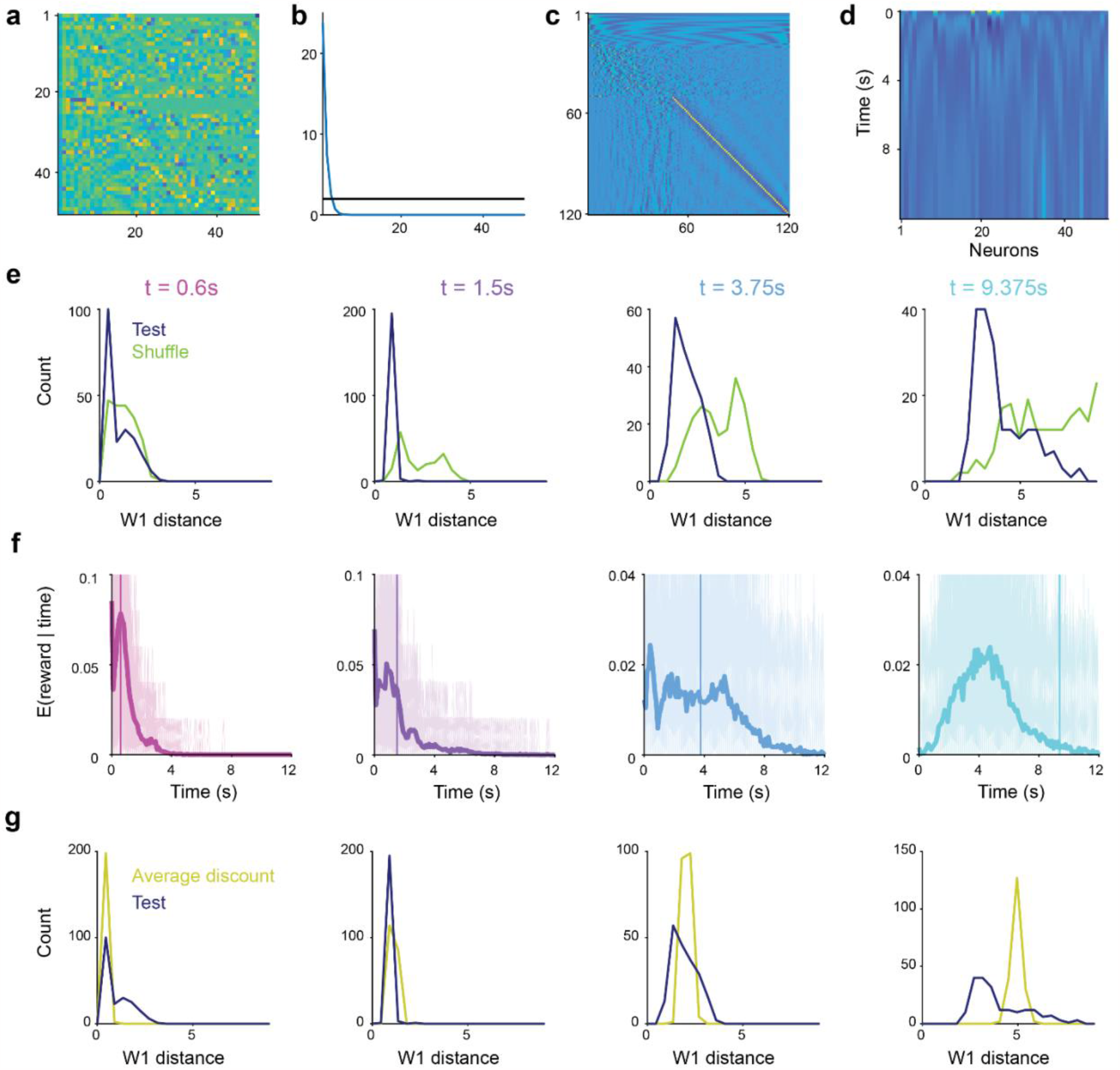
Decoding reward timing using the regularized pseudo-inverse of the discount matrix. **(a-c)**, Singular value decomposition (SVD) of the discount matrix. **a**, left singular vectors (in the neuron space). **b**, Singular values. The black line at 2 indicates the values of the regularization term α. **c**, right singular vectors (in the time space). **d**, Decoding matrix based on the regularized pseudo-inverse. **e**, Distribution of 1-Wassertein distances between the reward timing and the predicted reward timing from the decoding on the test data exponential fits (shown in Fig. 3k, top row) and on the shuffled data (shown if Fig. 3k, bottom row). The prediction from the test data are better predictions (smaller 1-Wasserstein distance) than shuffled data (*P* = 1.2 x 10^-4^ for 0.6 s reward delay, *P* < 1.0 x 10^-20^ for the other delays, one-tailed Wilcoxon signed rank test, see Methods). **f**, Decoded subjective expected timing of future reward *E*(*r*|*t*) using a model with a single discount factor (the mean discount factor across the population, see Methods). **g**, Distribution of 1-wassertein distances between the reward timing and the predicted reward timing from the decoding on the test data from exponential fits (shown in Fig. 3k, top row) and on the average exponential model (shown in **f**). Decoding is better for the exponential model from Fig. 3 than the average exponential model except for the shortest delay (*P*(*t* = 0.6s) = 1, *P*(*t* = 1.5s) < 1.0 x 10^-31^, *P*(*t* = 3.75) = 0.0135, *P*(*t* = 9.375s) < 1.0 x 10^-14^), one-tailed Wilcoxon signed rank test, see Methods).

**Extended Data Fig 6.**
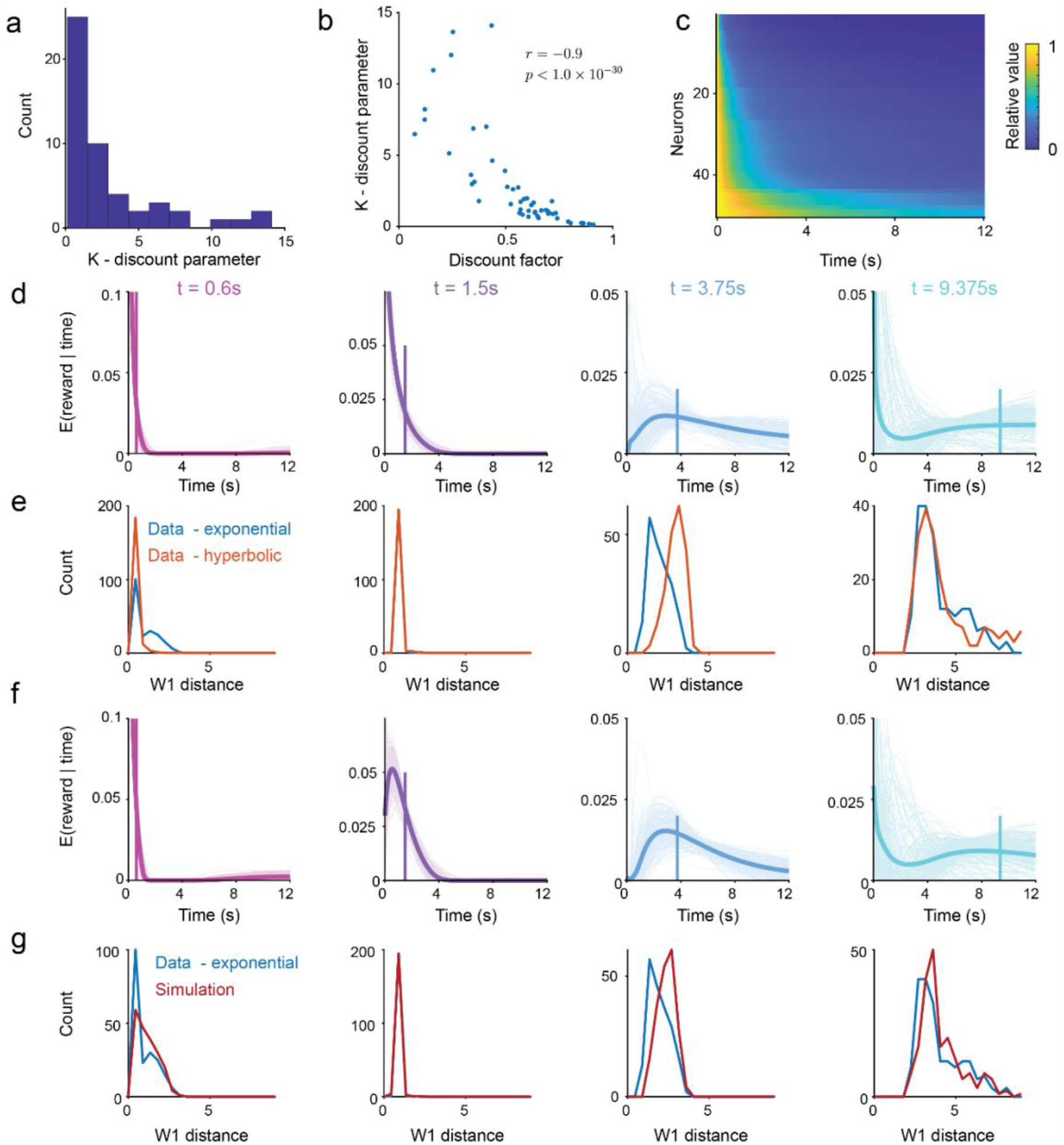
Decoding reward timing from the first to the hyperbolic model and exponential model simulations. **a**, Distribution of the inferred discount parameter k across the neurons. **b**, Correlation between the discount factor inferred in the exponential model of the discount parameter k from the hyperbolic model (*r* = -0.9, *P* < 1.0 x 10^-30^, *t*-test). Note the in the hyperbolic model a larger value of k implies faster discounting hence the negative correlation. **c**, Discount matrix for the hyperbolic model. For each neuron we plot the relative value of future events given its inferred discount parameter. Neurons are sorted by decreasing estimated value of the discount parameter. **d**, Decoded subjective expected timing of future reward *E*(*r*|*t*) using the discount matrix from the hyperbolic model (see Methods). **e**, Distribution of 1-Wassertein distances between the reward timing and the predicted reward timing from the decoding on the test data with the exponential model (shown in Fig. 3k, top row) and on the test data with the hyperbolic model (shown in **d**). Decoding is better for the exponential model from Fig. 3 than the hyperbolic model except for the shortest delay (*P*(*t* = 0.6s) = 1, *P*(*t* = 1.5s) < 1.0 x 10^-31^, *P*(*t* = 3.75) < 1.0 x 10^-33^, *P*(*t* = 9.375s) < 1.0 x 10^-3^), one-tailed Wilcoxon signed rank test, see Methods). **f**, Decoded subjective expected timing of future reward *E*(*r*|*t*) using simulated data based on the parameters of the exponential model (see Methods). **g**, Distribution of 1-Wassertein distances between the reward timing and the predicted reward timing from the decoding on the test data from exponential fits (shown in Fig. 3k, top row) and on the simulated data from the parameters of the exponential fits (shown in **f**). Decoding is marginally better for the data predictions (*P*(*t* = 0.6s) = 0.002, *P*(*t* = 1.5s) = 0.999, *P*(*t* = 3.75) <1 x 10^-12^, *P*(*t* = 9.375s) = 0.027), one-tailed Wilcoxon signed rank test, see Methods), suggesting that decoding accuracy is limited by the number of trials.

**Extended Data Fig 7.**
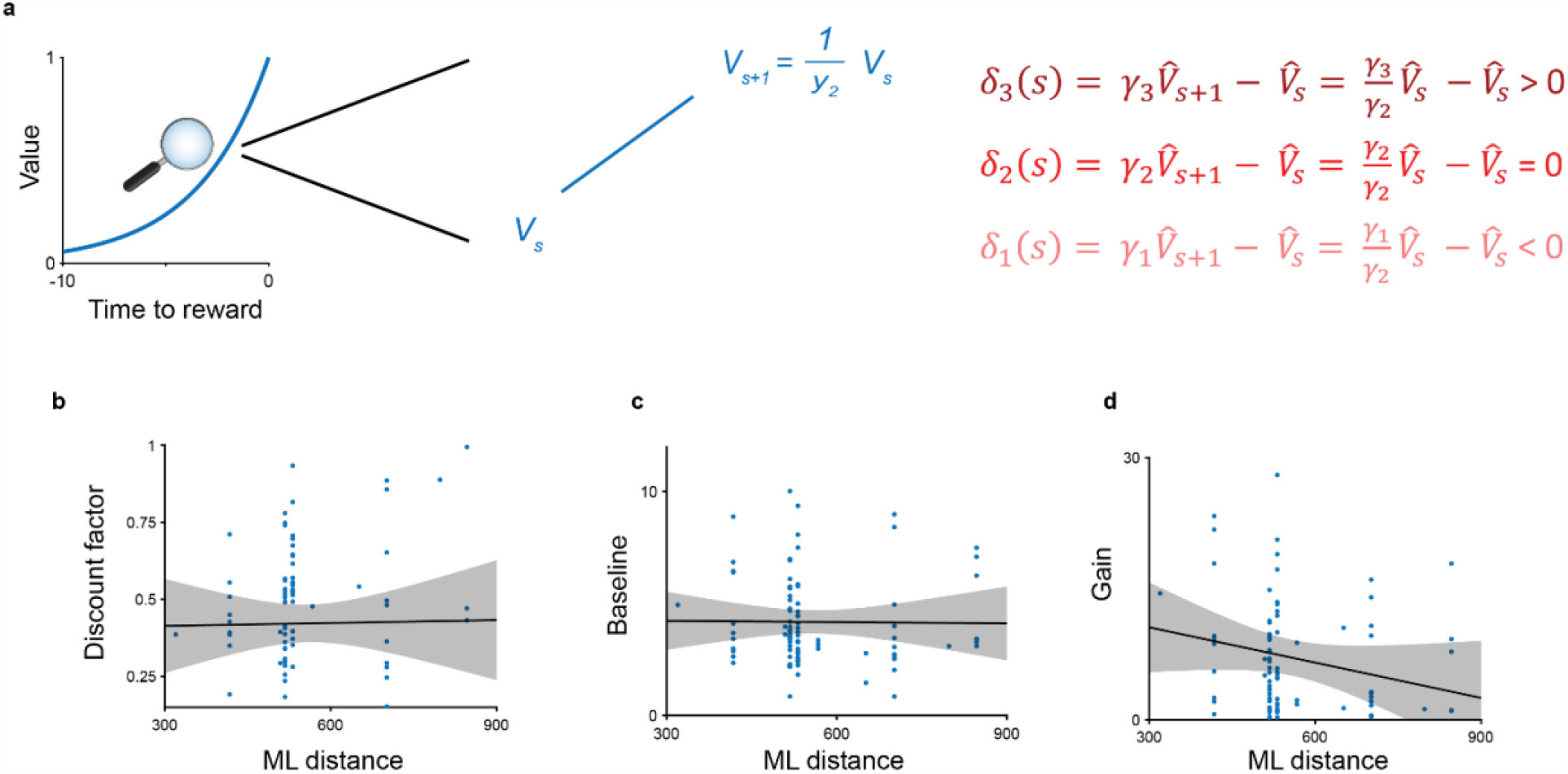
Ramping, discounting and anatomy. **a**, Ramping in the prediction error signal is controlled by the relative contribution of value increases and discounting. If the value increase (middle) exactly matches the discounting, there is no prediction error (middle equation, right). If the discounting is smaller than the value increase (large discount factor) then there is a positive TD error (top equation, right). If the discounting is larger (small discount factor) than the value increase then there a negative TD error (bottom equation, right). A single timescale agent with no state uncertainty will learn an exponential value function but if there is state uncertainty (see ref[^69^]) or the global value function arises from combining the contribution of single-timescale agents then the value function is likely t be non-exponential. **b**, The discount factor inferred in the VR task is not correlated with the medio-lateral (ML) position of the implant (Pearson’s *r* = 0.015, *P* = 0.89). **c**, The baseline parameter inferred in the VR task is not correlated with the medio-lateral (ML) position of the implant (Pearson’s *r* = -0.011, *P* = 0.92). **d**, The inferred gain in the VR task reduces with increasing medio-lateral (ML) position but the effect does not reach significance (Pearson’s *r* = -0.19, *P* = 0.069).

**Extended Data Fig 8.**
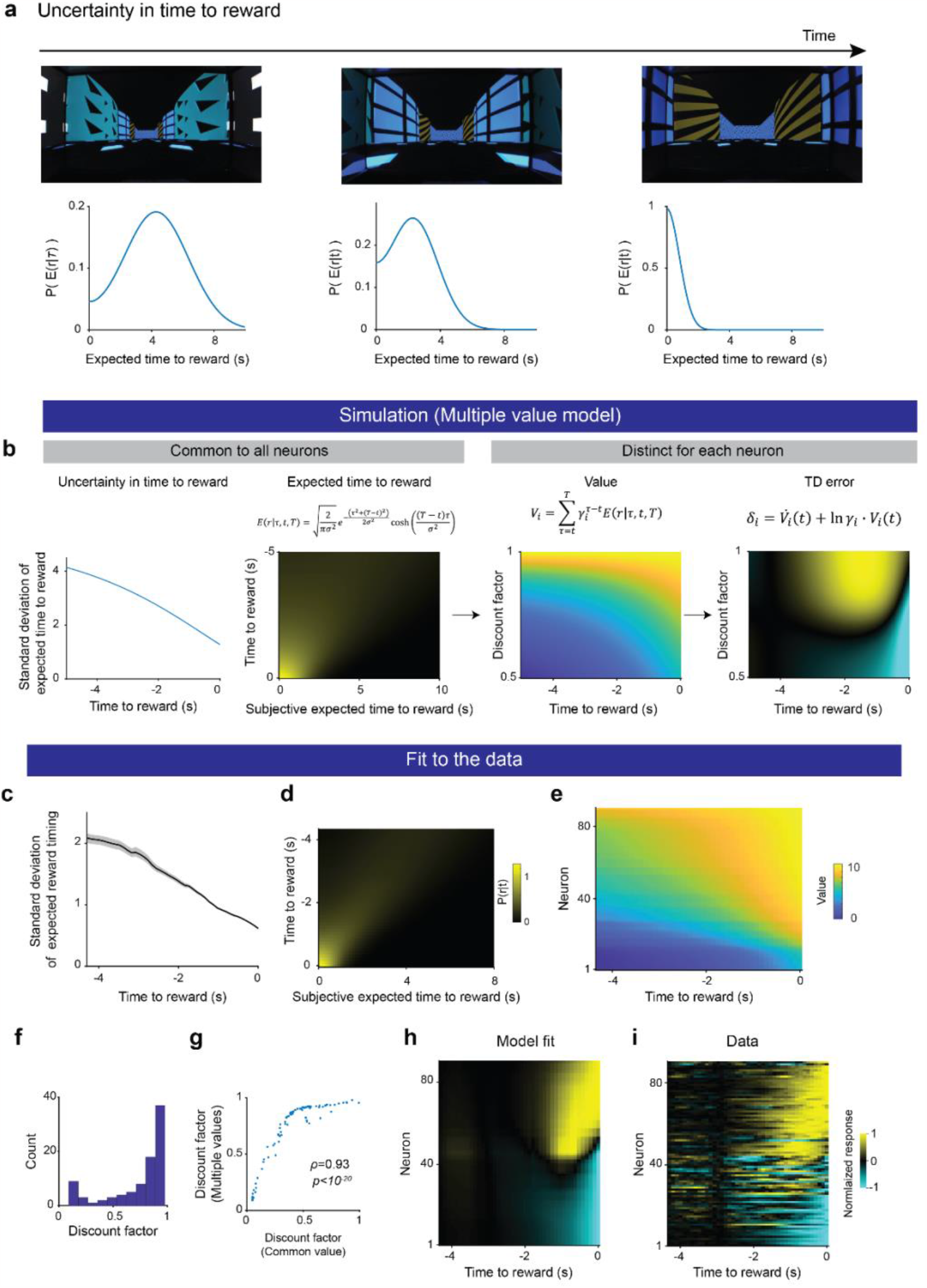
Discounting heterogeneity explains ramping diversity in a common reward expectation model. **a**, Uncertainty in reward timing reduces as mice approach the reward zone. Not only does the mean expected reward time reduces but the standard deviation of the estimate also reduces. Distribution in the bottom row from fitted data (see panels **c-i**). **b**, Simulations showing how reduction in uncertainty in reward timing (shared across neurons) and diverse discount factors lead to heterogeneous ramping activity in dopamine neurons. First panel. In this model, the uncertainty in the subjective estimate of reward timing (measured by the standard deviation) reduces as the mice approach the reward. Second panel. Distribution of subjective expected time to reward as a function of the true time to reward. The distribution is sampled from a folded normal distribution. The standard deviation reduces as reward approaches as shown in the first panel. Third panel. Given the subjective expected time to reward, common to all neurons due to a single world mode, we can compute a value function for each neuron given its discount factor. Fourth panel. This leads to a heterogeneity of TD errors across neurons, including monotonic upward and downwards ramps as well as non-monotonic ramps. **c**, The inferred standard deviation of the reward expectation model reduces as a function of time to reward. Line indicates the mean inferred standard deviation and the shading indicates the standard error of the mean over 100 bootstraps. **d**, Subjective expected timing of the reward as a function of true time to reward. As the mice approach the reward not only does the mean expected time to reward reduces but the uncertainty of the reward timing captured by the standard deviation shown in **c** also reduces. This effect leads to increasingly convex value functions that lead to the observed ramps in dopamine neuron activity. **e**, Value function for each individual neuron. **f**, Distribution of inferred discount factors under the common reward expectation model. g, Although the range of discount factor between the fits from the common value (x-axis) and common reward expectation (y-axis) models differs, the inferred discount factors are strongly correlated for single neurons (Spearman’s *ρ* = 0.93, *P* < 1.0 x 10^-20^). **h**, Predicted ramping activity from the model fits under the common reward expectation model. **i**, Diversity of ramping activity across single neurons as mice approach reward (aligned by inferred discount factor in the common reward expectation model).

**Extended Data Fig 9.**
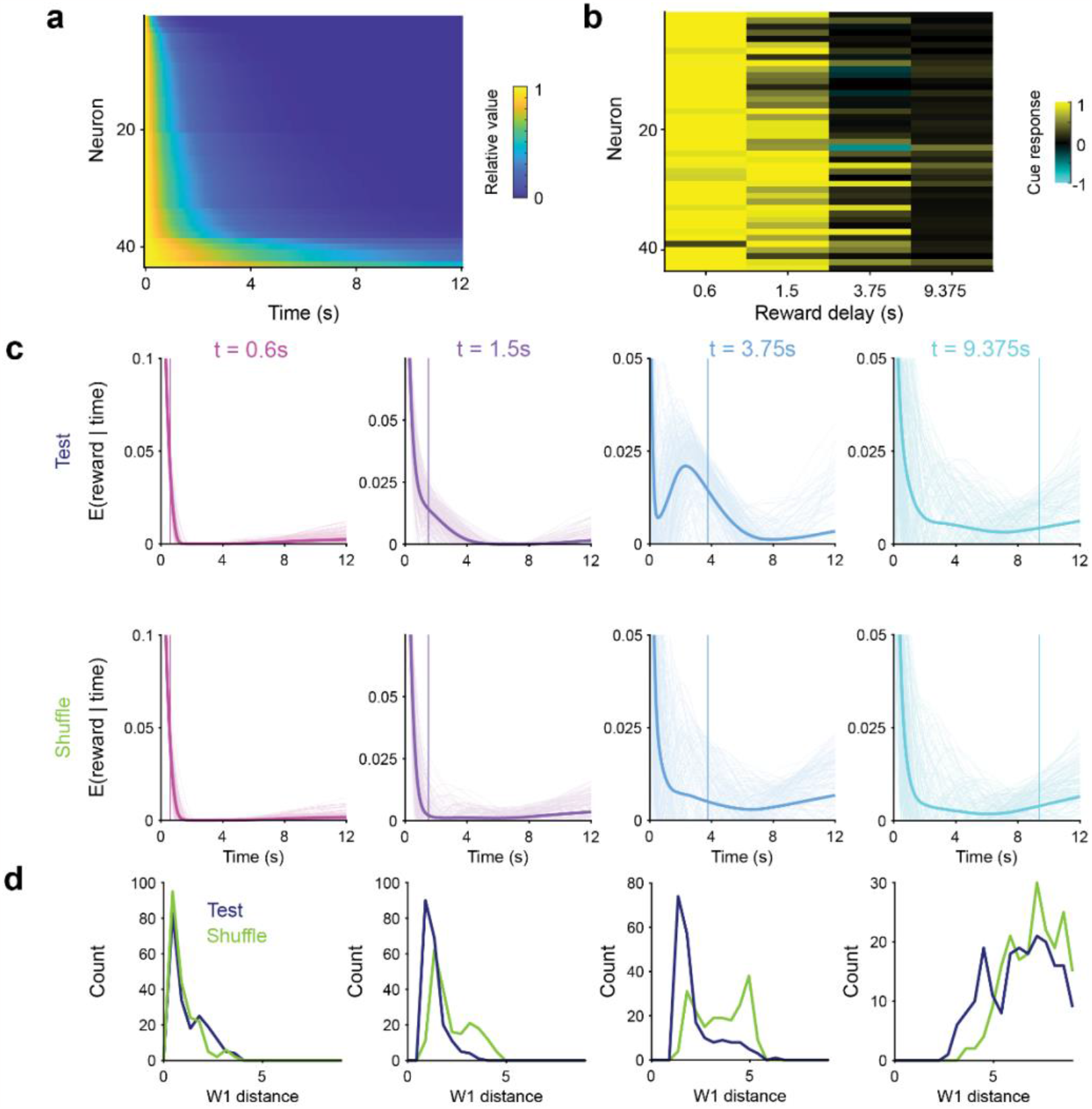
Decoding reward timing in the cud delayed reward task using parameters inferred in the VR task. **a**, Discount matrix computed using the parameters inferred in the VR tasks for neurons recorded across both tasks and used in the cross-task decoding. **b**, Dopamine neurons cue responses in the cued delay task. Neurons are aligned as in **a** according to increasing discount factor inferred in the VR task. **c**, Top row: Decoded reward timing using discount factors inferred in the VR task. Bottom row: The ability to decode reward timing is lost when shuffling the identities of the cue responses. **d**, Except for the shortest delay, decoded reward timing is more accurate than shuffle as measured by the 1-Wassertsein distance (*P*_*t* = 0.6s_ = 1, *P*_*t* = 1.5s_ < 1.1 x 10^-20^, *P*_*t* = 3.75s_ < 3.8 x 10^-20^, *P*_*t* = 9.375s_ < 2.9 x 10^-5^).

## References

1. Sutton, R. S. & Barto, A. G. Reinforcement Learning: An Introduction (Adaptive Computation and Machine Learning series). 552 (A Bradford Book, 2018).

2. Tesauro, G. Temporal difference learning and TD-Gammon. Commun. ACM 38, 58–68 (1995).

3. Mnih, V. et al. Human-level control through deep reinforcement learning. Nature 518, 529–533 (2015).

4. Silver, D. et al. Mastering the game of Go with deep neural networks and tree search. Nature 529, 484–489 (2016).

5. Ecoffet, A., Huizinga, J., Lehman, J., Stanley, K. O. & Clune, J. First return, then explore. Nature 590, 580–586 (2021).

6. Wurman, P. R. et al. Outracing champion Gran Turismo drivers with deep reinforcement learning. Nature 602, 223–228 (2022).

7. Schultz, W., Dayan, P. & Montague, P. R. A neural substrate of prediction and reward. Science 275, 1593–1599 (1997).

8. Schultz, W. Neuronal reward and decision signals: from theories to data. Physiol. Rev. 95, 853–951 (2015).

9. Cohen, J. Y., Haesler, S., Vong, L., Lowell, B. B. & Uchida, N. Neuron-type-specific signals for reward and punishment in the ventral tegmental area. Nature 482, 85–88 (2012).

10. Commons, M. L., Mazur, J. E., Nevin, J. A. & Rachlin, H. Effect Of Delay And Of Intervening Events On Reinforcement Value. 344 (Taylor & Francis Group, 2013).

11. Ainslie, G. Specious reward: a behavioral theory of impulsiveness and impulse control. Psychol. Bull. 82, 463–496 (1975).

12. Frederick, S., Loewenstein, G. & O’Donoghue, T. Time Discounting and Time Preference: A Critical Review. J. Econ. Lit. 40, 351–401 (2002).

13. Laibson, D. Golden Eggs and Hyperbolic Discounting. Q. J. Econ. 112, 443–478 (1997).

14. Sozou, P. D. On hyperbolic discounting and uncertain hazard rates. Proceedings of the Royal Society B: Biological Sciences 265, 2015–2020 (1998).

15. Rao, R. P. & Ballard, D. H. Predictive coding in the visual cortex: a functional interpretation of some extra-classical receptive-field effects. Nat. Neurosci. 2, 79–87 (1999).

16. Keller, G. B. & Mrsic-Flogel, T. D. Predictive processing: A canonical cortical computation. Neuron 100, 424–435 (2018).

17. LeCun, Y. A Path Towards Autonomous Machine Intelligence. https://openreview.net/forum?id=BZ5a1r-kVsf (2022).

18. Sutton, R. S., Bowling, M. H. & Pilarski, P. M. The Alberta Plan for AI Research. arXiv (2022) doi:10.48550/arxiv.2208.11173.

19. Sutton, R. S. Learning to predict by the methods of temporal differences. Mach. Learn. 3, 9–44 (1988).

20. Lillicrap, T. P. et al. Continuous control with deep reinforcement learning. arXiv (2015) doi:10.48550/arxiv.1509.02971.

21. Narasimhan, K., Kulkarni, T. & Barzilay, R. Language Understanding for Text-based Games Using Deep Reinforcement Learning. arXiv (2015) doi:10.48550/arxiv.1506.08941.

22. Mnih, V. et al. Asynchronous Methods for Deep Reinforcement Learning. arXiv (2016) doi:10.48550/arxiv.1602.01783.

23. Botvinick, M. et al. Reinforcement learning, fast and slow. Trends Cogn Sci (Regul Ed) 23, 408–422 (2019).

24. Gardner, M. P. H., Schoenbaum, G. & Gershman, S. J. Rethinking dopamine as generalized prediction error. Proc. Biol. Sci. 285, (2018).

25. Dabney, W. et al. A distributional code for value in dopamine-based reinforcement learning. Nature 577, 671–675 (2020).

26. Cox, J. & Witten, I. B. Striatal circuits for reward learning and decision-making. Nat. Rev. Neurosci. 20, 482–494 (2019).

27. Watabe-Uchida, M. & Uchida, N. Multiple dopamine systems: weal and woe of dopamine. Cold Spring Harb. Symp. Quant. Biol. 83, 83–95 (2018).

28. Engelhard, B. et al. Specialized coding of sensory, motor and cognitive variables in VTA dopamine neurons. Nature 570, 509–513 (2019).

29. Gershman, S. J. & Uchida, N. Believing in dopamine. Nat. Rev. Neurosci. 20, 703–714 (2019).

30. Hamid, A. A. et al. Mesolimbic dopamine signals the value of work. Nat. Neurosci. 19, 117–126 (2016).

31. Mohebi, A. et al. Dissociable dopamine dynamics for learning and motivation. Nature 570, 65–70 (2019).

32. Berke, J. D. What does dopamine mean? Nat. Neurosci. 21, 787–793 (2018).

33. Ainslie, G. Specious reward: a behavioral theory of impulsiveness and impulse control. Psychol. Bull. 82, 463–496 (1975).

34. Dasgupta, P. & Maskin, E. Uncertainty and hyperbolic discounting. American Economic Review 95, 1290–1299 (2005).

35. Dasgupta, P. Discounting climate change. J. Risk Uncertain. 37, 141–169 (2008).

36. Redish, A. D. Addiction as a computational process gone awry. Science 306, 1944–1947 (2004).

37. Milenkova, M. et al. Intertemporal choice in Parkinson’s disease. Mov. Disord. 26, 2004–2010 (2011).

38. Lempert, K. M. & Phelps, E. A. The malleability of intertemporal choice. Trends Cogn Sci (Regul Ed) 20, 64–74 (2016).

39. Lempert, K. M., Steinglass, J. E., Pinto, A., Kable, J. W. & Simpson, H. B. Can delay discounting deliver on the promise of RDoC? Psychol. Med. 49, 190–199 (2019).

40. Sutton, R. S. et al. Horde: A Scalable Real-Time Architecture for Learning Knowledge from Unsupervised Sensorimotor Interaction. in The 10th International Conference on Autonomous Agents and Multiagent Systems - Volume 2 761–768 (International Foundation for Autonomous Agents and Multiagent Systems, 2011).

41. Bellemare, M. G., Dabney, W. & Rowland, M. Distributional reinforcement learning. (The MIT Press, 2023).

42. Stadie, B. C., Levine, S. & Abbeel, P. Incentivizing Exploration In Reinforcement Learning With Deep Predictive Models. arXiv (2015) doi:10.48550/arxiv.1507.00814.

43. Schaul, T., Quan, J., Antonoglou, I. & Silver, D. Prioritized Experience Replay. arXiv (2015) doi:10.48550/arxiv.1511.05952.

44. Jaderberg, M. et al. Reinforcement Learning with Unsupervised Auxiliary Tasks. arXiv (2016) doi:10.48550/arxiv.1611.05397.

45. Tano, P., Dayan, P. & Pouget, A. A local temporal difference code for distributional reinforcement learning. NeurIPS 33, 13662–13673 (2020).

46. Brunec, I. K. & Momennejad, I. Predictive representations in hippocampal and prefrontal hierarchies. J. Neurosci. 42, 299–312 (2022).

47. Mauk, M. D. & Buonomano, D. V. The neural basis of temporal processing. Annu. Rev. Neurosci. 27, 307–340 (2004).

48. Buhusi, C. V. & Meck, W. H. What makes us tick? Functional and neural mechanisms of interval timing. Nat. Rev. Neurosci. 6, 755–765 (2005).

49. Tsao, A., Yousefzadeh, S. A., Meck, W. H., Moser, M.-B. & Moser, E. I. The neural bases for timing of durations. Nat. Rev. Neurosci. 23, 646–665 (2022).

50. Fiorillo, C. D., Newsome, W. T. & Schultz, W. The temporal precision of reward prediction in dopamine neurons. Nat. Neurosci. 11, 966–973 (2008).

51. Mello, G. B. M., Soares, S. & Paton, J. J. A scalable population code for time in the striatum. Curr. Biol. 25, 1113–1122 (2015).

52. Soares, S., Atallah, B. V. & Paton, J. J. Midbrain dopamine neurons control judgment of time. Science 354, 1273–1277 (2016).

53. Enomoto, K., Matsumoto, N., Inokawa, H., Kimura, M. & Yamada, H. Topographic distinction in long-term value signals between presumed dopamine neurons and presumed striatal projection neurons in behaving monkeys. Sci. Rep. 10, 8912 (2020).

54. Kiebel, S. J., Daunizeau, J. & Friston, K. J. A hierarchy of time-scales and the brain. PLoS Comput. Biol. 4, e1000209 (2008).

55. Kurth-Nelson, Z. & Redish, A. D. Temporal-difference reinforcement learning with distributed representations. PLoS ONE 4, e7362 (2009).

56. Botvinick, M. M., Niv, Y. & Barto, A. C. Hierarchically organized behavior and its neural foundations: a reinforcement learning perspective. Cognition 113, 262–280 (2009).

57. Shankar, K. H. & Howard, M. W. A scale-invariant internal representation of time. Neural Comput. 24, 134–193 (2012).

58. Tanaka, S. C. et al. Prediction of immediate and future rewards differentially recruits cortico-basal ganglia loops. Nat. Neurosci. 7, 887–893 (2004).

59. Wei, W., Mohebi, A. & Berke, J. Striatal dopamine pulses follow a temporal discounting spectrum. BioRxiv (2021) doi:10.1101/2021.10.31.466705.

60. Fedus, W., Gelada, C., Bengio, Y., Bellemare, M. G. & Larochelle, H. Hyperbolic Discounting and Learning over Multiple Horizons. arXiv (2019).

61. Sherstan, C., Dohare, S., MacGlashan, J., Günther, J. & Pilarski, P. M. Gamma-Nets: Generalizing Value Estimation over Timescale. AAAI 34, 5717–5725 (2020).

62. Momennejad, I. & Howard, M. W. Predicting the future with multi-scale successor representations. BioRxiv (2018) doi:10.1101/449470.

63. Kobayashi, S. & Schultz, W. Influence of reward delays on responses of dopamine neurons. J. Neurosci. 28, 7837–7846 (2008).

64. Schultz, W. Dopamine reward prediction-error signalling: a two-component response. Nat. Rev. Neurosci. 17, 183–195 (2016).

65. Matsumoto, H., Tian, J., Uchida, N. & Watabe-Uchida, M. Midbrain dopamine neurons signal aversion in a reward-context-dependent manner. eLife 5, (2016).

66. Howe, M. W., Tierney, P. L., Sandberg, S. G., Phillips, P. E. M. & Graybiel, A. M. Prolonged dopamine signalling in striatum signals proximity and value of distant rewards. Nature 500, 575–579 (2013).

67. Gershman, S. J. Dopamine ramps are a consequence of reward prediction errors. Neural Comput. 26, 467–471 (2014).

68. Kim, H. R. et al. A Unified Framework for Dopamine Signals across Timescales. Cell 183, 1600–1616.e25 (2020).

69. Mikhael, J. G., Kim, H. R., Uchida, N. & Gershman, S. J. The role of state uncertainty in the dynamics of dopamine. Curr. Biol. 32, 1077–1087.e9 (2022).

70. Guru, A. et al. Ramping activity in midbrain dopamine neurons signifies the use of a cognitive map. BioRxiv (2020) doi:10.1101/2020.05.21.108886.

71. Doya, K. Reinforcement learning in continuous time and space. Neural Comput. 12, 219–245 (2000).

72. Lee, R. S., Engelhard, B., Witten, I. B. & Daw, N. D. A vector reward prediction error model explains dopaminergic heterogeneity. BioRxiv (2022) doi:10.1101/2022.02.28.482379.

73. Lowet, A. S., Zheng, Q., Matias, S., Drugowitsch, J. & Uchida, N. Distributional reinforcement learning in the brain. Trends Neurosci. 43, 980–997 (2020).

74. Millidge, B. G., Song, Y., Lak, A., Walton, M. E. & Bogacz, R. Reward-Bases: Dopaminergic Mechanisms for Adaptive Acquisition of Multiple Reward Types. BioRxiv (2023) doi:10.1101/2023.05.09.540067.

75. Cruz, B. F. et al. Action suppression reveals opponent parallel control via striatal circuits. Nature 607, 521–526 (2022).

76. Eshel, N., Tian, J., Bukwich, M. & Uchida, N. Dopamine neurons share common response function for reward prediction error. Nat. Neurosci. 19, 479–486 (2016).

77. Menegas, W., Akiti, K., Amo, R., Uchida, N. & Watabe-Uchida, M. Dopamine neurons projecting to the posterior striatum reinforce avoidance of threatening stimuli. Nat. Neurosci. 21, 1421–1430 (2018).

78. Collins, A. L. & Saunders, B. T. Heterogeneity in striatal dopamine circuits: Form and function in dynamic reward seeking. J. Neurosci. Res. 98, 1046–1069 (2020).

79. Louie, K. Asymmetric and adaptive reward coding via normalized reinforcement learning. PLoS Comput. Biol. 18, e1010350 (2022).

80. Xu, Z., van Hasselt, H. P. & Silver, D. Meta-Gradient Reinforcement Learning. Advances in Neural Information Processing Systems (2018).

81. Yoshida, N., Uchibe, E. & Doya, K. Reinforcement learning with state-dependent discount factor. in 2013 IEEE Third Joint International Conference on Development and Learning and Epigenetic Robotics (ICDL) 1–6 (IEEE, 2013). doi:10.1109/DevLrn.2013.6652533.

82. Schlegel, M. et al. General value function networks. jair 70, 497–543 (2021).

83. Kvitsiani, D. et al. Distinct behavioural and network correlates of two interneuron types in prefrontal cortex. Nature 498, 363–366 (2013).

84. Arulkumaran, K., Deisenroth, M. P., Brundage, M. & Bharath, A. A. Deep reinforcement learning: A brief survey. IEEE Signal Process. Mag. 34, 26–38 (2017).

85. Oppenheim, A., Willsky, A. & Hamid, W. Signals and Systems. 1000 (Pearson, 1996).

86. Dayan, P. Improving Generalization for Temporal Difference Learning: The Successor Representation. Neural Comput. 5, 613–624 (1993).

87. Gershman, S. J. The successor representation: its computational logic and neural substrates. J. Neurosci. 38, 7193–7200 (2018).

88. Amit, R., Meir, R. & Ciosek, K. Discount Factor as a Regularizer in Reinforcement Learning. in (PMLR, 2020).

89. Badia, A. P. et al. Agent57: Outperforming the Atari Human Benchmark. in (PMLR, 2020).

90. Leone, F. C., Nelson, L. S. & Nottingham, R. B. The folded normal distribution. Technometrics 3, 543 (1961).

